# Preclinical antiviral study of a liver-targeted TLR1/2 agonist in an immune-competent mouse model of HBV infection

**DOI:** 10.64898/2026.05.27.728102

**Authors:** Fanny Charriaud, Myriam Lamrayah, Romain Barnault, Svenja Schuehle, Manon Desmares, Mathias Heikenwalder, Julie Lucifora, Bernard Verrier, David Durantel

**Affiliations:** Laboratoire de Biologie Tissulaire et d’Ingénierie Thérapeutique, CNRS_UMR5305, Université Claude Bernard Lyon 1 (UCBL1), Lyon, France; Centre International de Recherche en Infectiologie (CIRI), INSERM_U1111, CNRS_UMR5308, ENS-Lyon, Université Claude Bernard Lyon 1 (UCBL1), Lyon, France; Division of Chronic Inflammation and Cancer, German Cancer Research Center (DKFZ), Heidelberg, Germany

**Keywords:** Hepatitis B virus, Preclinical study, TLR1/2-ligand/agonist, Nano-vectorization, HBV cure

## Abstract

Chronic hepatitis B cure requires the inactivation and/or elimination of covalently closed circular DNA (cccDNA), together with silencing of integrated viral genomes and restoration of HBV-specific immune responses. The TLR1/2 agonist Pam_3_CSK_4_ has previously been identified as a potent direct anti-HBV agent *in vitro*. In the present study, we engineered a liver-targeting polymeric nanoparticle formulation of Pam_3_CSK_4_ to enhance its *in vivo* immunostimulatory and antiviral activity. We evaluated the antiviral efficacy of this novel nanoformulation carrying the TLR1/2 agonist (NP-Pam_3_CSK_4_) in monotherapy and started to investigate its mechanism of action through immunological correlates in an immune-competent AAV-HBV mouse model. AAV-HBV-infected mice received intravenous administrations of NP-Pam_3_CSK_4_ at doses of 5 or 20 μg twice per treatment cycle over four cycles, followed by a 2-week follow-up period. Soluble Pam_3_CSK_4_ was administered at substantially higher doses (100 μg). Serial blood samples were regularly collected to monitor virological and host immune parameters. At study completion, liver tissues were harvested for intrahepatic quantification of viral and immunological markers using immunoassays, quantitative PCR, and histological analyses. The most pronounced antiviral effects were observed in mice treated with NP-Pam_3_CSK_4_ formulations, which achieved greater viral suppression than free Pam_3_CSK_4_ despite markedly lower administered doses. Histological examination of liver biopsies from treated animals revealed prominent immune cell infiltration, including macrophages, monocytes, and T cells, organized in dense cluster-like structures. These findings support the induction of coordinated innate and adaptive immune responses contributing to HBV control and clearance. Collectively, our results demonstrate that nanoparticle-based delivery of TLR1/2 agonist represents a promising therapeutic strategy for chronic HBV infection and may improve the likelihood of achieving functional cure. Further mechanistic and translational studies (combination) are warranted to support clinical development.

## Introduction

Food and Drug Administration-approved treatments for chronic Hepatitis B infection (CHB) include nucleos(t)ide analogues (NUCs) and pegylated-interferon-alfa (Peg-IFN-a) (European Association for the Study of the Liver, 2025). While NUCs are highly effective at suppressing viral replication by targeting the HBV polymerase and exhibit an excellent safety profile, they rarely induce hepatitis B surface antigen (HBsAg) seroclearance, a currently desired therapeutic endpoint. Furthermore, long-term (often long-life) administration is generally required, as treatment discontinuation or arrest is associated with near-universal viral rebound. In contrast, Peg-IFN-a, either alone or in combination with NUCs, leads to higher rates of HBsAg loss (Zhang & Terrault, 2024), despite limited tolerability due to frequent side effects. These observations indicate that an innate immune stimulation might represent an important therapeutic component, which could leverage multiple mode of action for achieving HBV cure (Dusheiko et al., 2023; Gehring & Protzer, 2019; Lucifora et al., 2014; Zhao et al., 2024). In line with this statement, several drugs are currently undergoing phase III clinical trials for CHB, following successful phase II assessments, are mainly antisense oligonucleosides (*e. g*. bepirovirsen, AHB-137), which are also endowed with toll-like-receptor (TLR) stimulation capacities (*i. e*. mainly TLR8) (Bouquet et al ., 2026; Hong & Rajwanshi, 2025). Additional drugs acting as TLR or RIG1/NOD2-like agonists, including TLR7 (*e. g*. GS-9620/vesatolimod) and TLR8 (*e. g*. GS-9688/selgantolimod), were/are being investigated in phase II clinical trials (Boni et al., 2018; E. Gane et al., 2021; E. J. Gane et al., 2023; Hu et al., 2021; H. L. A. Janssen et al., 2018; Yuen et al., 2023). While these agents have shown limited efficacy as monotherapies, they are now further considered in combination strategies. If the RIG/NOD-like receptor agonist, Inarigivir Soproxil, caused one death during trials, none of the TLR agonists/ligands administrated to human so far have caused lethal/detrimental adverse effect, leaving avenues for the improvement for these types of therapeutic components.

One such avenue is the use of TLR agonists/ligands capable to directly inhibit the replication of HBV, in addition to their indirect effect in restoring immune response. Our recent studies demonstrated that TLR1/2 agonists, unlike TLR7 and TLR8 agonists, are capable to directly inhibit HBV replication in hepatocytes (*i. e*. primary human hepatocytes and differentiated HepaRG cells). Mechanistically, TLR1/2 activation induces the activation of NF-κB pathways, leading to a strong reduction of HBV replication intermediates in these *in vitro* models (Desmares et al., 2022; Lucifora et al ., 2018; Michelet et al., 2022).

The present study participates in efforts to optimize and validate a novel TLR1/2-based therapeutic component capable of both directly inhibiting HBV replication in hepatocytes and inducing immune-mediated HBV control. Such dual activity may impact on the breadth of HBV replication, reduce the number of viral episome-containing cells, and increase rates of long-term HBsAg clearance. Indeed, studies highlighted that TLR2 ligands significantly increase cellular immunity against HBV, and that TLR2-deficient mice have deficient HBV-specific CD8+ T cells (Dou et al., 2020; Ma et al., 2017; Naghib et al., 2022). Collectively, these findings suggest that TLR1/2 agonists may provide a broader antiviral profile than TLR7 or TLR8-targeted approaches. Yet, our previous work indicated that intravenous (IV) administration of soluble Pam_3_CSK_4_ (escalating doses from 20 to 80 μg/IV/mouse) in HBV-infected liver-humanized mice achieved only limited antiviral efficacy (Lucifora et al., 2018). To increase therapeutic efficacy, we developed a novel Pam_3_CSK_4_ formulation by chemically optimizing its incorporation into polylactic acid (PLA)-based nanoparticles (NP) and demonstrated preservation of both structural and functional integrity of the resulting nano-vectors (NP-Pam_3_CSK_4_) (Lamrayah et al., 2019, 2023). Here, we evaluated the antiviral efficacy of NP-Pam_3_CSK_4_ monotherapy *in vivo* and initiated investigations on immune correlates associated with antiviral activity. Experiments were conducted using a preclinical HBV-replicating mouse model, based on adeno-associated virus (AAV-HBV) transduction (Dion et al., 2013), which features high level of viral episomes (both AAV-HBV and *bona fide* cccDNA) (Frederick, 1987; Lucifora et al ., 2017). This model mimics an “immune-tolerant” phenotype of chronic HBV infection, relevant for evaluating immunomodulatory antiviral strategies.

## Material and methods

### Cell lines and Culture

HepaRG cells (wild type (wt) and TLR2 or TLR3 knocked-out (KO)) were cultured and infected with HBV as previously described (Desmares et al., 2022; Lucifora et al., 2018).

### *Chemical synthesis of NP, fluorescent NP and NP-Pam*_**3**_***CSK***_**4**_

PLA NP were prepared by nanoprecipitation as previously described (Lamalle-Bernard et al., 2006). Briefly, the polymer was dissolved in acetone and added dropwise to an aqueous solution under stirring. Organic solvents were then removed under reduced pressure at 30 °C (rotavapor R-300; Buchi). Either Pam_3_CSK_4_ (InvivoGen, USA) or the near-infrared fluorescent probe DiR XenoLight Dye (λ_excitation_ = 750 nm, λ_emission_ = 770 nm) (Perkin Elmer, USA) was incorporated during the nanoprecipitation. Precisely, Pam_3_CSK_4_ was incorporated following the published procedure with a drug:PLA ratio of either 0.26 % or 0.80 % (Lamrayah et al., 2019). The final Pam_3_CSK_4_ concentration in the colloidal suspension was 35 µg/mL (low concentration) or 130 µg/mL (high concentration) for a PLA concentration of 15 mg/mL. Adjuvatis SA (France) manufactured the near-infrared DiR XenoLight Dye NP with a fluorophore:PLA ratio of 0.02 %. The average hydrodynamic diameter and size distribution (mentioned as polydispersity index) of formulations (n=7/condition) were determined by dynamic light scattering at 25 °C and a scattering angle of 173° using a Zetasizer Nano ZS (Malvern, UK). The colloidal suspensions were diluted in 1 mM NaCl solution, and each value was the mean of four independent measurements. The electrophoretic mobilities (mentioned as zeta potential) were measured by laser Doppler velocimetry using the same equipment, at a scattering angle of 12.5°.

### Animal-scale imaging of fluorescent NP delivery

Ten-week-old SKH1 mice (n=4/experiment) were purchased from Charles River Laboratories (France) and acclimated for 7 days. All animals were kept under specific pathogen-free (SPF) conditions in the P-PAC animal platform (Lyon, France) and experimental protocol s undertaken were approved by the ethical committee of the facility (ministerial authorization number 2017022719265782). At week-11, animals received retro-orbital IV injections of fluorescent NP (100 µL, 0.96 µM of fluorescent probe/injection). On the day of imaging, mice were anesthetized under 4 % isoflurane. Live measurement and monitoring of the fluorescence were made using either a near-infrared open imaging system (Kaer Imaging System, KAER Labs, France) or a fluorescence molecular tomography system (2D/3D FMT4000, Perkin Elmer, USA). For the latter, the collected fluorescence data were reconstructed by TrueQuant software (Perkin Elmer, USA) for the quantification of three-dimensional fluorescence signals.

### AAV-HBV persistent infection mouse model

Five-week-old female C57BL/6 mice were purchased from Charles River Laboratories (France) and acclimated for 7 days. All animals were kept under SPF conditions in A3 animal facility (PBES, Lyon, France) and experimental protocols undertaken were approved by the ethical committee of Ecole Normale Supérieure de Lyon (ministerial authorization number 37673-2022040520076454). Six-weeks-old mice received an hepatotropic recombinant AAV 2/8 vector carrying 1.3 genome of HBV-genotype D, serotype ayw (rAAV2/8-1.3HBV, 10^11^ vge (virus genome equivalent)/mice, 100 μL in PBS) (Dion et al., 2013) by retro-orbital intravenous (IV) injection. The transduction went on for 35 days to launch HBV replication. Four days before the onset of treatment (at day-31), animals were bled and allocated into groups (n=7/group) with comparable viremia and HBsAg levels.

### Experimental design

The *in vivo* study was divided into four treatment cycles of 10 days each. One treatment cycle consisted of two IV injections and one blood sampling, resulting to a total of 8 injections/animal. All injections were made retro-orbitally following anesthesia (150 µL, 4 % isoflurane). The control group was treated with 3TC (Epivir 10mg/ml, ViiV Healthcare), administered in drinking water (at 100 mg/kg/day) and protected from light. Non-infected and non-treated groups received PBS 1X by IV. Soluble Pam_3_CSK_4_ group received 100 µg/injection. NP-Pam_3_CSK_4_ low and NP-Pam_3_CSK_4_ high groups received the equivalent of respectively 5 µg and 20 µg of vectorized Pam_3_CSK_4_. At day-74, viremia was assessed to determine whether a fourth treatment cycle was required; persistent viral replication warranted administration of such a fourth cycle and pursue the experiment until day-90. Blood samples (days 42, 53, 64, 74 and 90) were collected retro-orbitally to monitor serum viremia and antigenemia overtime. At endpoint (day-90), liver and blood were collected for further analysis. Experiments were performed during the light phase of the day. Mice weight was monitored every week, and all efforts were made to minimize suffering.

### Blood analysis

Serum was collected after heating fresh blood at 37 °C for 30 minutes, followed by two centrifugations (10 minutes, 16000 xg), and stored at -80 °C. HBsAg levels in the serum (at 100-, 1000- and 10000-fold dilutions) were determined by a chemiluminescence immunoassay kit (CLIA) according to manufacturer’s instructions (Autobio, China). CLIA plates were analyzed using the Luminoskan^™^ (ThermoFisher, France). HBV DNA copies were measured by using real-time quantitative PCR (qPCR) after nucleic acid extraction with Nucleospin RNA Virus kit according to the manufacturer’s instructions (Macherey-Nagel, Germany). Lactate deshydrogenase (LDH) levels were analyzed according to manufacturer’s instructions (LDH-Glo^™^ cytotoxicity assay, Promega, France) on diluted serum (100-fold).

### Intrahepatic RNA and DNA extraction from hepatocytes

Liver biopsies were snap-frozen in liquid nitrogen at endpoint and stored at -80 °C. Tissue was homogenized on ice in TE buffer using Dounce’s glass (Sigma Aldrich, USA). Homogenized tissue was then separated in two fractions for cell dissociation with either TRI Reagent® (Zymo Research, USA) for total RNA or with TE Buffer-10 % SDS for total DNA in the presence of boiled RNAse A and proteinase K. Total RNA was further isolated by chloroform extraction and isopropanol precipitation, while total DNA was isolated with phenol/chloroform/isoamyl alcohol phase separation and ethanol precipitated in presence of glycogen. Finally, total RNA was dissolved in RNAse-free water and total DNA was dissolved in TE buffer. Nucleic acids were quantified by NanoDrop^™^ (ThermoFisher, France) and stored at -80 °C at 500 ng/µL.

### Reverse transcription and qPCR analysis

Total RNA treated with RQ1 DNase (Promega, France) was reverse transcribed using the Maxima First Strand synthesis Kit as instructed by provider (K1641, ThermoFisher Scientific, USA). Complementary DNA was stored at -80 °C at 10 ng/µL. To evaluate the level of intrahepatic HBV episomes (cccDNA + AAV-HBV), a T5 exonuclease digestion was performed as instructed by provider (New England Biolabs, UK), and as described in (Lucifora et al., 2017). Real-time qPCR reactions were performed with either Sybr Green mix Luna® 2X (New England Biolabs, UK) or TaqMan^™^ fast advanced 1X (Applied Biosystems, USA) with a 10 µL total qPCR mix and 4 µL DNA or cDNA input. Primers used are listed in **Table 1**. Reactions were performed using an Applied QuantStudio^™^ 1 machine. For viremia, absolute quantification was performed with pregenomics (pg) RNA primers. For intrahepatic measures, DNA or mRNA expression levels were quantified using the comparative cycle threshold (Ct) method. Relative quantification was performed using the ΔΔCt method, normalizing to housekeeping gene (GAPDH or TFRC) and PBS-treated mice.

**Table 1.**
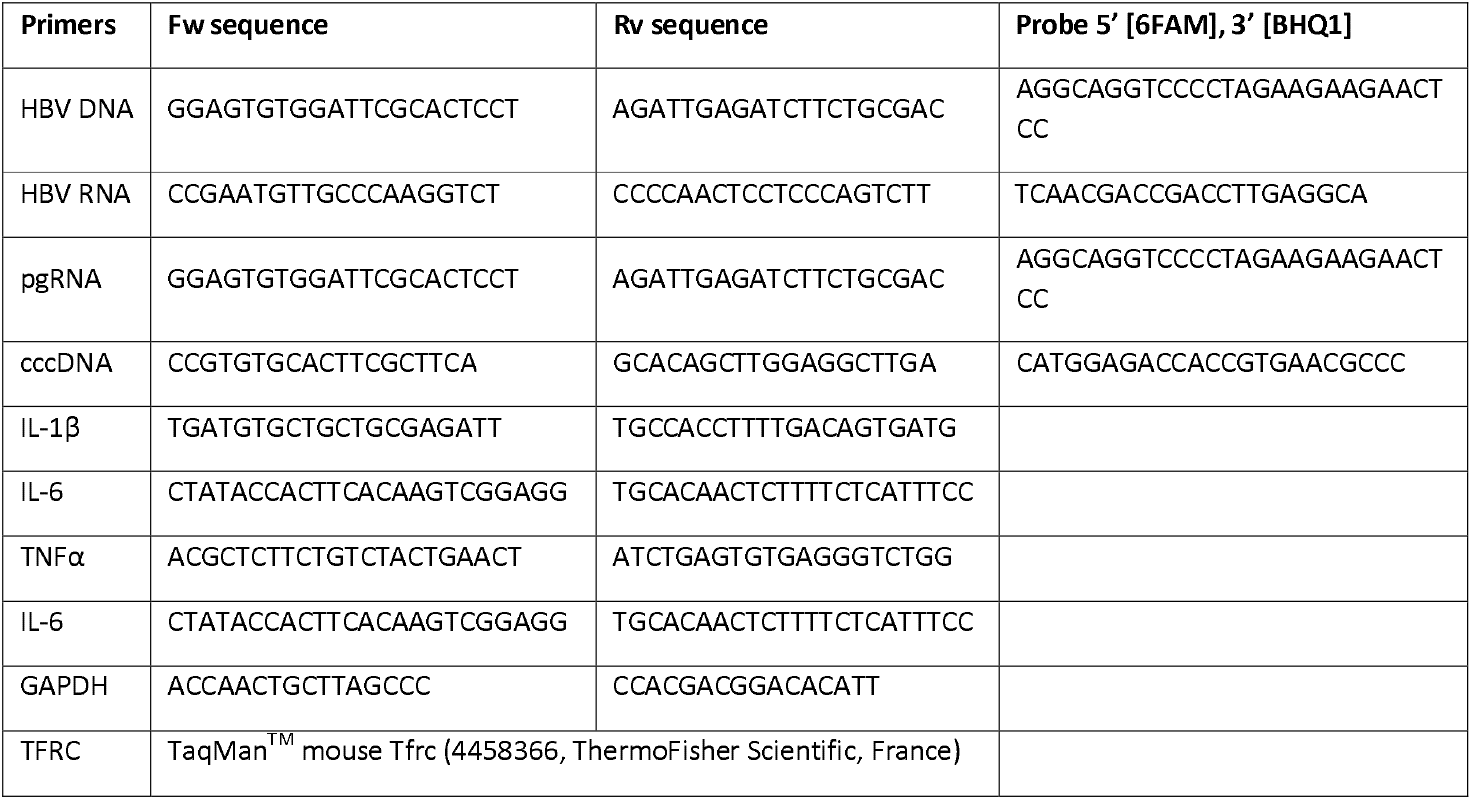

### Liver hematoxylin/eosin (H&E) staining and automated cluster quantification

At endpoint, liver tissue pieces (2/animal) were fixed in 4 % buffered formalin for 24 hours at 4 °C and paraffin embedded. Then, 4.5-μm-thin liver sections were prepared with a rotary microtome (CM1860 Ag protect, LEICA, Germany) and mounted on slides. Slides were deparaffinized, rinsed, stained with hematoxylin (5 minutes) and then counter stained with eosin (5 minutes) for structural assessment of the tissue integrity. Brightfield images were automatically acquired overnight using the ZEISS Axio scan.Z1 slide scanner (Zeiss, Germany) available at the CIQLE facility (LyMic Lyon Multiscale Imaging Center). The x40 magnification (numerical aperture 0.95) was used for all acquisitions, with a pixel height and width of 0.1100 µm. For image segmentation and quantification of cell clusters, QuPath version 0.4.2 was used (Bankhead et al., 2017). More precisely, a pixel classifier was trained with a set of five images to distinguish background and portal areas versus (vs) organ tissue. A second pixel classifier was trained with a set of eight images to distinguish clusters vs normal tissue. The StarDist whole-cell segmentation deep learning-based method was used for nuclear anatomical detection (Schmidt et al., 2018). An all-in-one script was applied to all images before cell abundance and cluster quantification were extracted from QuPath for graphical representation.

### Liver immunostaining

For immunohistochemical (IHC) staining, liver pieces were fixed in ROTI®HistoFix 4 % (Carl Roth) for 48h, which was then changed to 70 % ethanol. Tissues were placed into embedding cassettes and IHC was performed as previously described (Wolf et al., 2014). Briefly, 2 µm sections of formalin-fixed, paraffin-embedded livers were cut and then stained with indicated antibodies using a Leica BOND-MAX. Antibodies used are listed in **Table 2**. Slides were scanned on a SCN400 slide scanner (Leica) and analyzed using ImageJ software.

**Table 2.**

| Antigen | Dilution | Provider | Catalog number |
| --- | --- | --- | --- |
| CD3 | 1/500 | InvitroGen | MA1-90582 |
| CD8 | 1/400 | Cell signaling | 98941S |
| CD4 | 1/1000 | ThermoFisher | 14-9766-82 |
| MHCII | 1/500 | Novus Biologicals | NBP1-43312 |
| Clec4f | 1/1000 | R&D | AF2784-SP |
| F4/80 | 1/250 | Linaris | T-2006 |
| CD11b | 1/10000 | Abcam | Ab133357 808 |
| HBc | 1/100 | Origen | AP10430PU-N |
| CD206 | 1/500 | Abcam | Ab64693 |

### Statistical analysis

Experimental data were reported as mean + SD. Statistical analyses were performed using GraphPad Prism software v8 (GraphPad, USA). Normality or lognormality was assessed according to Shapiro-Wilk test. If positive, one-way or two-way ANOVA with Dunnett’s multiple comparison to PBS group was performed. If not, Kruskal-Wallis test with Dunnett’s multiple comparisons to PBS group was performed. Mann-Whitney test was also used for some analysis. Threshold for statistical significance was 0.05 for all analysis, with *p < 0.05, **p < 0.01, and ***p < 0.005 ; ns means non-significative.

## Results

### PLA-formulated TLR1/2 agonist maintains its anti-HBV activity in vitro

We previously reported the anti-HBV activity of non-vectorized Pam_3_CSK_4_ *in vitro* using HBV-infected differentiated-HepaRG (dHepaRG) cells (Desmares et al., 2022; Lucifora et al., 2018). Using the same model and experimental protocol (**Fig. S1A**), we investigated whether the novel formulation of Pam_3_CSK_4_ will affect its antiviral activity and specificity profile. Results show NP-Pam_3_CSK_4_ formulation was as efficient as Pam_3_CSK_4_ to reduce levels of HBV intracellular RNAs (**Fig. S1B**) and secreted antigens (**Fig. S1C**), without any toxicity (up to 1500 ng/mL (equivalent NP numbers were used to compare empty NP vs Pam_3_CSK_4_ loaded NP); **Fig. S1D**). The Efficient Concentration 50 % (EC_50_) of NP-Pam_3_CSK_4_ was at around 60 ng/mL (*i. e*. for intracellular HBV RNA parameter), which is slightly lower than the 15 ng/mL reported in *Desmares and colleagues* work (Desmares et al., 2022). Importantly, results obtained in TLR2- and TLR3-KO HepaRG cell lines demonstrated the antiviral activity of NP-Pam_3_CSK_4_ depends on TLR2 engagement (**Fig. S1E**), thus indicating that Pam_3_CSK_4_ loading into PLA-NP did not change its specificity of action through TLR1/2-MDA5-NF-κB axis. All these *in vitro* criteria validated the use of NP-Pam_3_CSK_4_ *in vivo*.

### Tissue-scale distribution analysis shows direct access and accumulation of PLA-NP in the liver of mice

Although PLA-NP were previously used by us (Lamrayah et al., 2019, 2023) and others (Peres et al., 2017; Shakya et al., 2023) to improve the drug delivery to various organs, no extensive study of their liver retention upon IV injection was performed to date. Fluorescent PLA-NP were retro-orbitally IV injected in mice and whole-body live imaging showed an immediate liver accumulation with a residual signal maintained for up to two weeks (**Fig. S2A/B**). More precisely, the liver delivery was already observed after only 15 seconds and a lobe segmentation analysis at 360 seconds indicated the homogeneous distribution across all liver lobes (**Fig. S2C**). As expected, one-hour post-injection PLA-NP seemed to be mainly taken up by macrophages (**Fig. S2D**). Altogether, these data suggest the potential for prolonged intrahepatic immune stimulation by NP-Pam_3_CSK_4_, and inform the design of the *in vivo* administration schedule used in subsequent cohort studies (**Fig. 1**).

**Figure 1.**
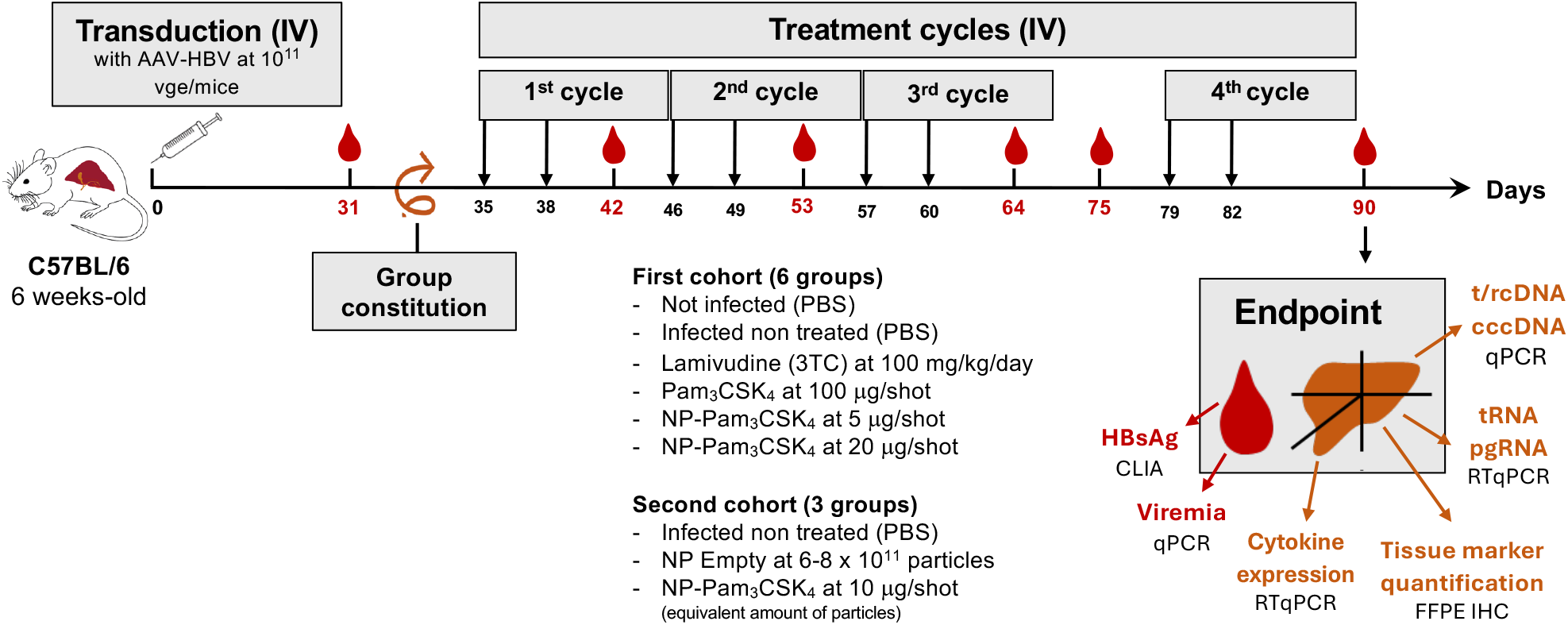
Cohort study design. Six-weeks-old C57BL/6 male mice were mock-transduced or transduced with 10^11^⍰vge of AAV-HBV 1.3 mer genotype D/serotype ayw for 31 days to allow establishment of chronic infection. At day-35, treatment cycles were started, after mice were evenly grouped according to viremia and HBsAg antigenemia. Mice (n⍰= ⍰7/group) were injected, at indicated days with vehicle (PBS), free/non vectorized Pam_3_CSK_4_ at 100 μg/injection, NP-Pam_3_CSK_4_ at 5 μg/shot, or NP-Pam_3_CSK_4_ at 20 μg/shot. 3TC was administrated continously at approximately 100 mg/kg/day (in drinking water) between days-35 and 64 and days-79 and 90. Blood/serum was collected at indicated times (blood drop symbol). Mice were euthanized at 90 days post-transduction. The different readouts and used techniques for exploitation of blood and liver samples are indicated.

### Antiviral effect of NP-Pam_3_CSK_4_ on HBV viremia and HBsAg antigenemia in AAV-HBV transduced C57BL/6 mice

An AAV-HBV transduction mouse model was used, as initially described (Dion et al ., 2013) to establish persistent HBV replication through hepatocytes delivery of high viral genome loads (10^11^ vge/mouse). HBV replication progressively increased and reached a plateau by day 35 post-transduction. When optimized in immune-competent systems (such as C57BL/6 mice), AAV-HBV transduction leads to a persistent and robust accumulation of intrahepatic HBV RNAs, transcribed from both AAV-HBV-episomes and *bona fide* cccDNA (Lucifora et al., 2017), formed by recombination according to previous work (Ko et al., 2021). This is accompanied by intrahepatic HBV proteins, as well as a high circulating levels of HBV virions (viremia > 10^7^ copies of genome/mL) and HBsAg subviral particles (>10^4^ IU/mL). Yet because murine hepatocytes do not express human NTCP, the HBV entry receptor, viral spread to naïve hepatocytes cannot occur. Despite this limitation, this model has been extensively used in antiviral and vaccine studies (Herschke et al., 2021; Kosinska et al., 2019; Martin et al., 2015; Sacherl et al., 2023; Wang et al., 2023; Xu et al., 2023), and is widely recognized as a relevant preclinical model to evaluate immune therapies aimed at breaking HBV immune tolerance.

Our NP vectorized TLR1/2-ligand was tested under two conditions, which were compared side-by-side; 5 µg (NP-Pam_3_CSK_4_ low) or 20 µg (NP-Pam_3_CSK_4_ high) of nano-vectorized Pam_3_CSK_4_ vs 100 µg of free-form Pam_3_CSK_4_. *In vivo* injectable NP batches were quality-controlled; gold-standard colloidal properties of NP-Pam_3_CSK_4_ low and high formulations were found to have similar characteristics with a size of approximately 160 nm, a homogeneous distribution of the batches in suspension (polydispersity index < 0.100) and a similar negative surface charge (-70 mv) (**Fig. S3**).

Six-weeks-old mice received 10^11^ genome-particles of AAV-HBV and HBV replication was left establishing for 35 days before treatments. Between day-31 and day-35 post-transduction, 6 groups of 7 mice were formed by standardizing viremia and antigenemia. Then, four treatment cycles were performed (first dosing at day-35 and last dosing at day-82) with 2 IV injections per cycle (see detailed schedule in **Fig. 1**). A 10-day cycle corresponded to two retro-orbital IV injections plus one blood sampling, with eye alternance according to animal welfare recommendations. High HBV replication level was obtained in each group, with levels of viremia reaching in average 10^8^ copies/mL and corresponding HBsAg antigenemia at around 20 000 IU/mL at day-31 before the onset of treatment (*c. f*. homogenous baseline viremia and antigenemia in panels A of **Fig. 2 and 3**).

**Figure 2.**
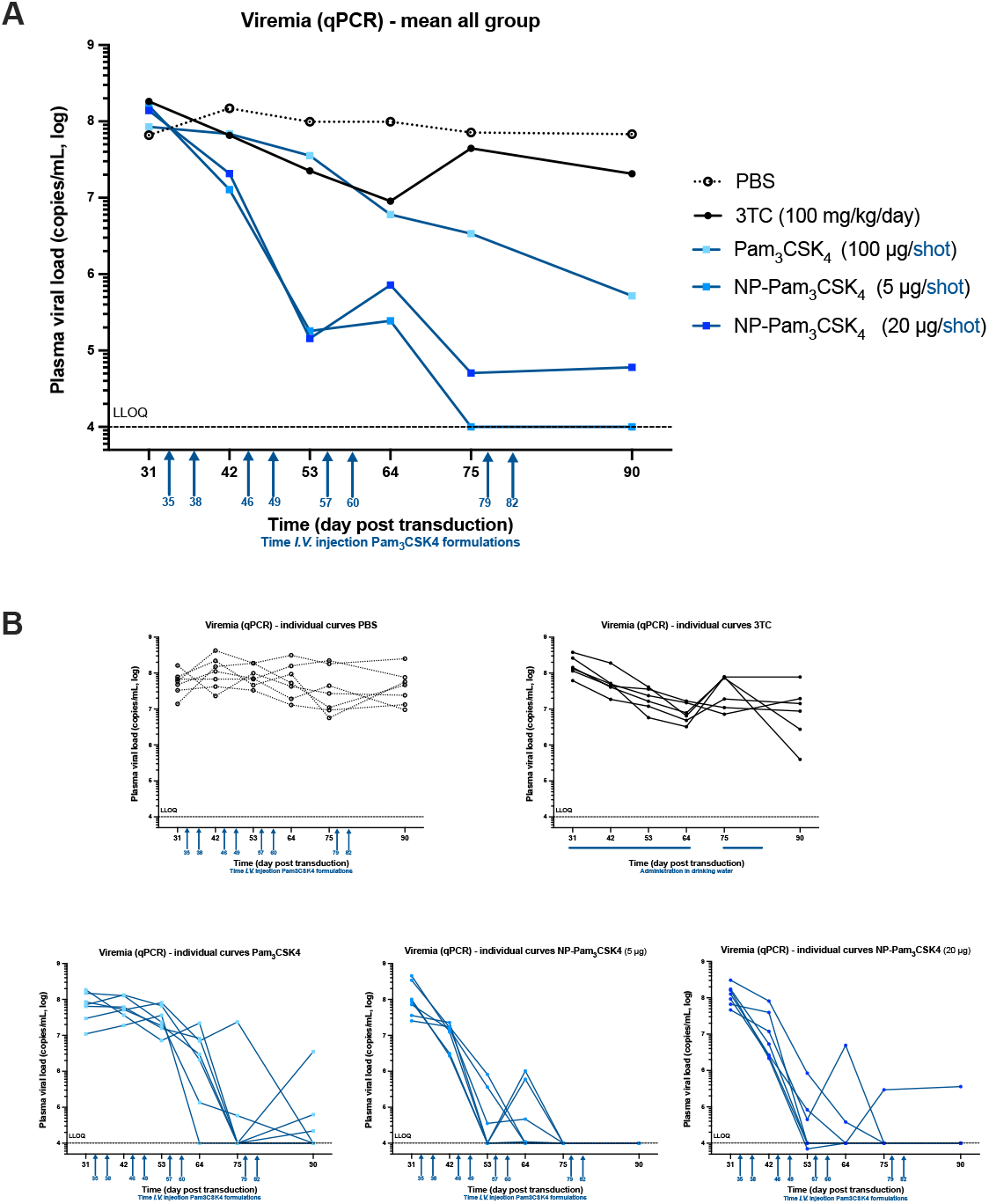
HBV viremia kinetics under various Pam_3_CSK_4_ treatments in infected mice. (A) HBV DNA was extracted from serum at days 31, 42, 53, 64, 75 and 90 post-transduction and subjected to qPCR. HBV standards were run in parallel to allow absolute quantification and presentation of results in copy/mL. Mean curves for each group are represented. (B) Viremia curves for each individual mouse, in each group, are represented.

**Figure 3.**
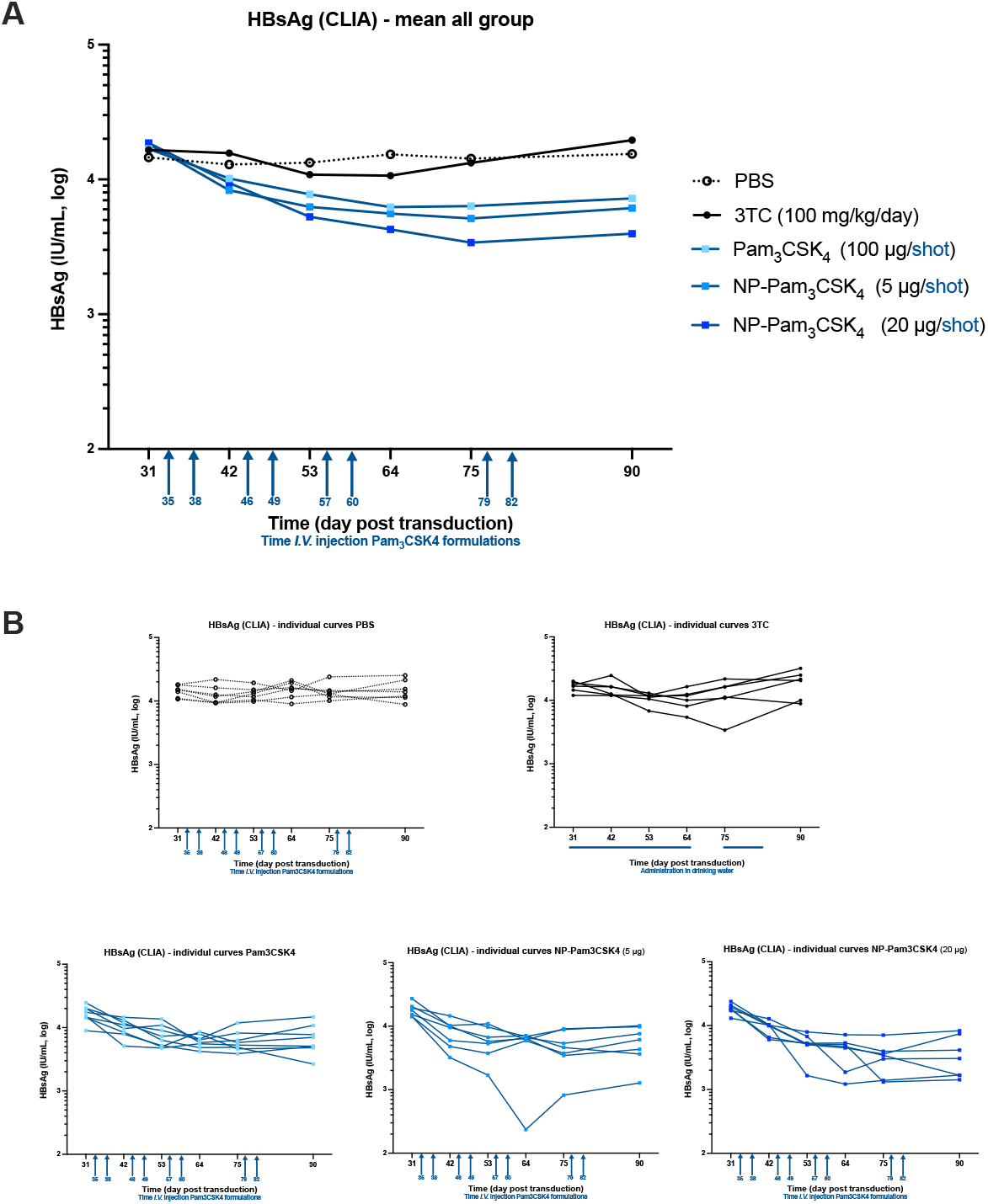
HBsAg antigenemia kinetics under various Pam_3_CSK_4_ treatments in infected mice. (A) HBsAg was quantified by CLIA at days 31, 42, 53, 64, 75 and 90 post-transduction. Three dilutions (1/100, 1/1000, and 1/10000) per condition were analyzed to allow technical robustness. Mean curves for each group are represented with HBsAg in IU/mL. (B) Antigenemia curves for each individual mouse, in each group, are represented.

Viremia analysis overtime highlighted a fast mean decrease of more than 3 log_10_ in NP-Pam_3_CSK_4_ high group at endpoint (*i. e*. euthanasia at day-90), and a > 4 log_10_ reduction, to undetectable level, in NP-Pam_3_CSK_4_ low group (**Fig. 2A and Fig. 4A**). In sharp contrast, in free Pam_3_CSK_4_ group, despite the respective 5-fold or 20-fold higher dose compared to NP-Pam_3_CSK_4_ groups, viremia was only reduced by around 2 log_10_ at day-90 (**Fig. 2A and 4A**). Of note, the first animals to reach undetectable viremia are in NP-Pam_3_CSK_4_ groups at day-53, whereas animal in Pam_3_CSK_4_ groups reached the same effect after the third treatment cycle only, at day-64 (one mouse) and day-75 (4 mice) (**Fig. 2B**). It is worth mentioning that all seven animals in the NP-Pam_3_CSK_4_ low group reached undetectability (under LLOQ) at the end of the experiment. It is also worth noting that 3TC, a nucleoside analogue used as control of anti-HBV activity, led to only a mild effect on viremia (around 1 log_10_ at day-64); this lack of efficacy was likely due to the route of administration in drinking water, which was not as efficient as oral gavage. As an “empty NP” control group was not included in the first cohort of mice, a second cohort was run with a similar protocol and 3 groups of 6 mice each, *i. e*. PBS, NP-empty, and NP-Pam_3_CSK_4_ (at 10 μg/injection). This second cohort confirmed the strong effect of NP-Pam_3_CSK_4_ on viremia and demonstrated that NP moiety alone does not likely contribute to the antiviral activity of NP-Pam_3_CSK_4_ (**Fig. S4**).

**Figure 4.**
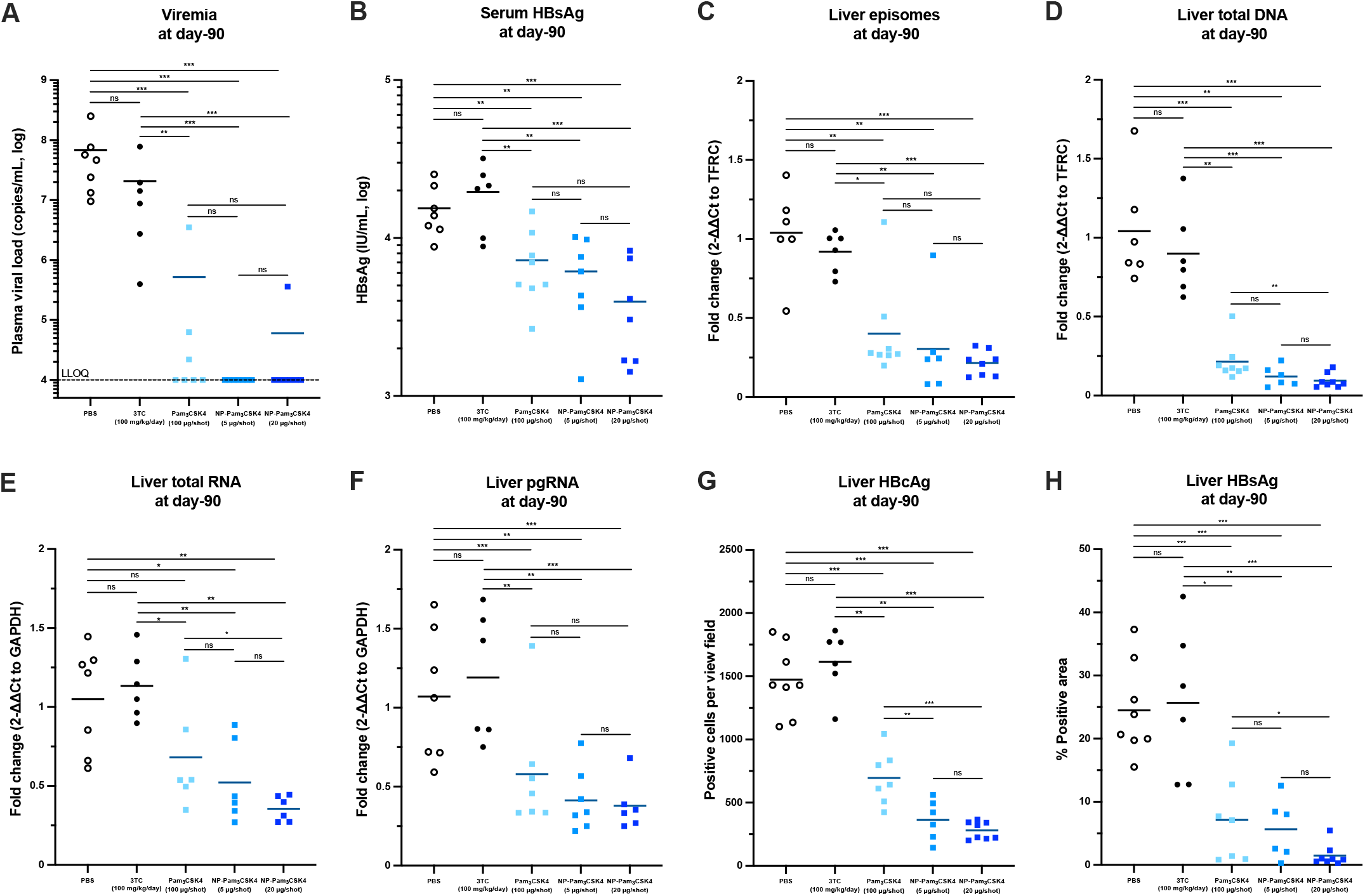
Virological analyses at euthanasia. At day-90, liver pieces were collected and either snap-frozen for nucleic acid extraction or fixed and paraffin embedded for IHC staining. (A) Viremia as analyzed in Figure 2 is represented at day-90. (B) HBsAg antigenemia as analyzed in Figure 3 is represented at day-90. (C) Intrahepatic DNAs were extracted/purified and digested with T5 exonuclease to remove excess of HBV rcDNA. Viral episomes (AAV-HBV + cccDNA) were quantified by qPCR with cccDNA specific primers. (D) DNAs were extracted and HBV total DNA (rcDNA mainly as episomes are negligible) quantified by qPCR. (E) Intrahepatic RNAs were extracted/purified and HBV total RNAs quantified by RTqPCR with primers located at end of HBV RNAs. (F) Intrahepatic RNAs were extracted/purified and HBV pgRNAs quantified by RTqPCR with primers located at 5’ of long HBV RNAs. (G) HBcAg and (H) HBsAg expression in hepatocytes were quantified following IHC staining. Threshold for statistical significance was 0.05 for all analysis, with *p < 0.05, **p < 0.01, and ***p < 0.005 ; ns means non-significative.

Albeit weaker than that seen for viremia, the most significative mean decrease in HBsAg levels was observed in NP-Pam_3_CSK_4_ high group, with an 82 % drop at day-75 (mean at 3 396 IU/mL) compared to day-31 (mean at 18 662 IU/mL) (**Fig. 3A**). At day-90 (endpoint), HBsAg mean level was respectively at 3 955 IU/mL (-76 %) in NP-Pam_3_CSK_4_ high group, 6 135 IU/mL (-67 %) in NP-Pam_3_CSK_4_ low group and 7 243 IU/mL (-61 %) in free-form Pam_3_CSK_4_ group, vs 15 455 IU/mL in PBS and 19 611 IU/mL in 3TC groups (**Fig. 4B**). There were no statistical differences between all Pam_3_CSK_4_ groups, but all Pam_3_CSK4 groups were statistically lower than control groups (**Fig. 4B**).

Treatments with NP-Pam_3_CSK_4_ led to a delayed mouse growth, which is not observed in free Pam_3_CSK_4_ group (**Fig. S5A**); however, at day-90 no significative changes in body weight could be evidenced between groups and explain the *in fine* antiviral effects. Such delay in body weight gain was nevertheless interpretated as a sign for compensated liver damaging. To explore the eventuality of the latter, LDH released in the blood from damaged cells was monitored overtime. All Pam_3_CSK_4_ treatments were associated to increased levels (2.5-fold at day-90) of LDH in serum (**Fig. S5B**), thus indicating some moderate hepatic toxicity and hepatocyte turnover. The latter was further confirmed my measurement of mean liver weights at euthanasia (1 g for non-infected group, 0.93 g for non-treated group, 0.94 g for 3TC group, 1.29 g for free Pam_3_CSK_4_ group, 1.13 g for NP-Pam_3_CSK_4_ low group, and 1.59 g for NP-Pam_3_CSK_4_ high group) and evidence of hepatomegaly in Pam_3_CSK_4_ treated groups.

### Intrahepatic virological analyses at endpoint indicates a variable effect on HBV replication in the different TLR1/2-stimulated groups

Next, intrahepatic analyses of HBV replication intermediates/parameters demonstrated that NP-Pam_3_CSK_4_ groups led to the strongest antiviral effect in mouse livers (**Fig. 4**), confirming serological analysis. Yet, no significative differences were observed between NP-Pam_3_CSK_4_ low (5 µg) and NP-Pam_3_CSK_4_ high (20 µg) groups for all parameters analyzed; only a trend toward a higher efficacy of the highest dose was observed. In details, we quantified viral episomes (AAV-HBV + cccDNA generated by homologous recombination, according to Ko and colleagues (Ko et al., 2021) by cccDNA-specific qPCR, total rcDNA by qPCR, total and pgRNAs by RTqPCR, and viral protein expressions (HBcAg and HBsAg) by immune-staining.

Effect on viral episomes ranged from 60 to 80 % reduction in Pam_3_CSK_4_ groups, with best effect obtained in NP-Pam_3_CSK_4_ high group (**Fig. 4C**). These reductions were significant when compared with PBS/3TC groups, but not significant between the different Pam_3_CSK_4_ groups. As expected 3TC did not affect viral episome level. In liver samples, this 3TC treatment did not lead either to a significative impact on total HBV DNA level (*i. e*. rcDNA), despite the fact nucleoside analogues are meant to inhibit HBV polymerase and decrease HBV rcDNA neosynthesis. The lack of activity of this control drug is likely due to a suboptimal administration route. In sharp contrast, Pam_3_CSK_4_ treatments, in particular with nano-particulated forms, led to a very significative decrease of intrahepatic HBV DNA (**Fig. 4D**), which was > 1 log_10_ in NP-Pam_3_CSK_4_ high group. These reductions in amount of total intrahepatic HBV DNA correlated well with results obtained for viremia and confirmed that the main effect of Pam_3_CSK_4_ agonists was on rcDNA neosynthesis, which in turn translates into a strong viremia reduction (> 3 log_10_ in NP-Pam_3_CSK_4_ groups).

Correlating respectively with viral episome and total HBV DNA (*i. e*. rcDNA) decreased levels, we also found that total and pgRNA levels were significantly decreased in Pam_3_CSK_4_ groups (**Fig. 4E and 4F**), with again the highest decrease observed in NP-Pam_3_CSK_4_ groups. As expected, due to the mode of action of this drug, no decrease of HBV RNA was observed in 3TC group; in fact, a trend toward an increased accumulation was noticed (**Fig. 4E and 4F**). These effects on HBV RNA accumulation in Pam_3_CSK_4_ groups translated into a reduction of intrahepatic HBV proteins as measured by quantification on slice staining (**Fig. S6**). Intrahepatic HBcAg staining was reduced by 75-80 % in NP-Pam_3_CSK_4_ groups, compared to PBS, and vs 50 % for free Pam_3_CSK_4_ (**Fig. 4G**). HBsAg-signal decrease in staining was particularly significant in NP-Pam_3_CSK_4_ high group (94 % decrease), in line with serological quantification of HBsAg by CLIA (**Fig. 4H**).

### Immune-staining and targeted gene expression correlate myeloid population infiltration/expansion with variable Pam_3_CSK_4_-induced antiviral activity

Based on numerous virological reads-out performed on serum and liver samples, our dataset has established a very potent anti-HBV effect of NP-Pam_3_CSK_4_ formulations. To get insights on immune driven antiviral mechanisms at work in Pam_3_CSK_4_-treated mice, liver tissue integrity and immune correlates of activity were explored based on IHC staining and targeted RTqPCR analysis.

Meshed amongst a healthy-looking tissue environment (absence of necrosis or damage), immune cell clusterization was noticeable only in the livers of Pam_3_CSK_4_ (*i. e*. TLR1/2-ligand) treated mice, suggesting immune cell proliferation and/or differentiation in these groups (**Fig. 5 and S7**). Observation of the tissue morphology suggests accumulation of these immune cells around sinusoids and portal triads (**Fig. 5A**). In comparison, PBS- and 3TC-treated mice did not show any signs of immune cell clusterization. Cell abundance was proportional to the antiviral effect for all Pam_3_CSK_4_ groups, with a highest score for NP-Pam_3_ CSK_4_ high group (94 clusters/4 mm^2^), followed by NP-Pam_3_ CSK_4_ low group (73 clusters/4 mm^2^) and nearly half as much for free form Pam_3_ CSK_4_ (50 clusters/4 mm^2^) (**Fig. 5B**). Similarly, total clustered area was the highest in NP-Pam_3_CSK_4_ high group (**Fig. 5C**).

**Figure 5.**
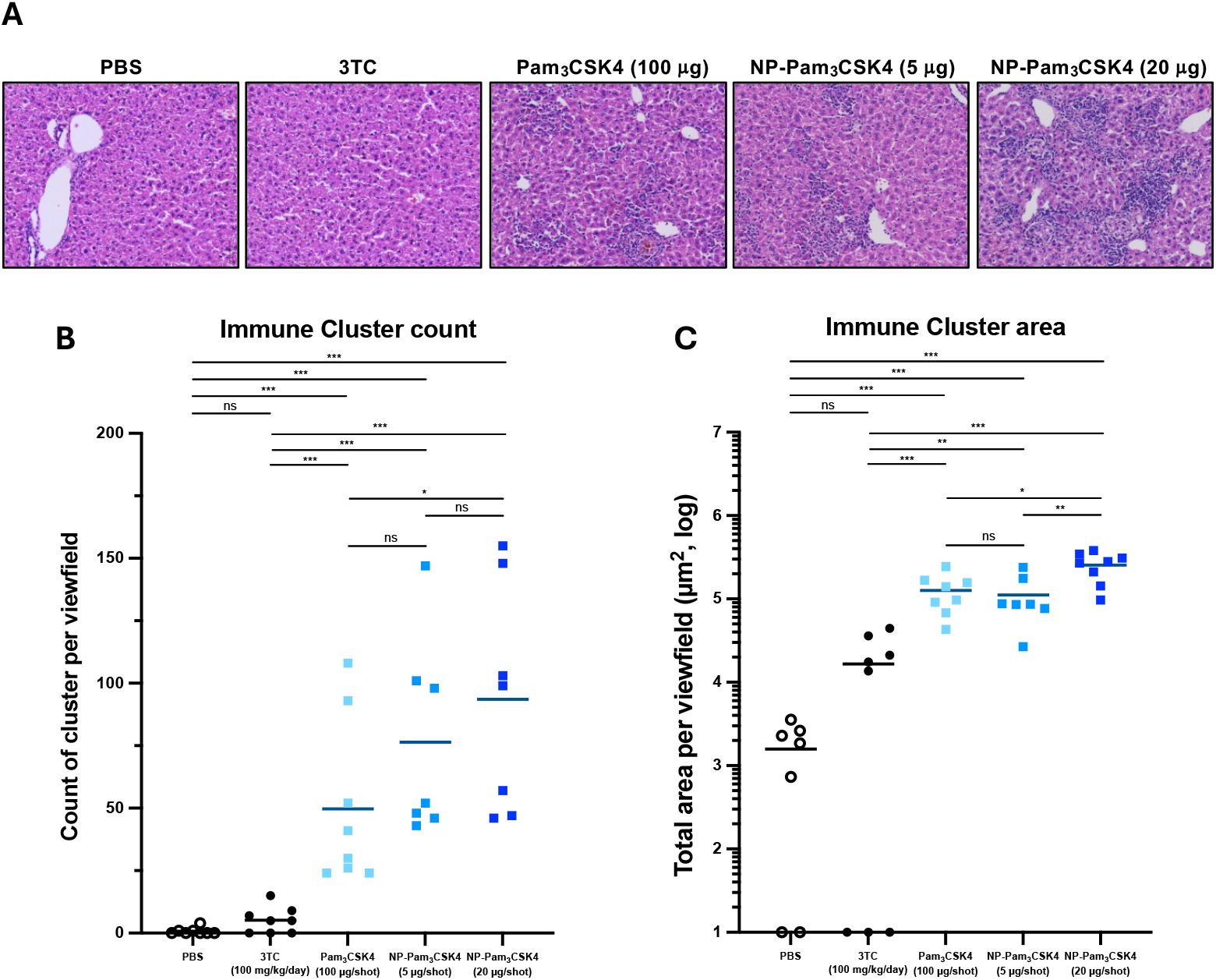
Quantification of immune cell clusters in liver parenchyma of Pam_3_CSK_4_-treated mice. (A) H&E staining was performed on fixed and paraffin-embedded liver slices. Brightfield images were automatically acquired using the ZEISS Axio scan.Z1 slide scanner. Quantification of cell clusters was done using QuPath version 0.4.2. An all-in-one script was applied to all images before cell abundance and cluster quantification were extracted from QuPath for graphical representation. (B) Immune cluster count. (C) Immune cluster area quantification. Threshold for statistical significance was 0.05 for all analysis, with *p < 0.05, **p < 0.01, and ***p < 0.005 ; ns means non-significative.

As TLR1/2 ligands target in particular macrophages/monocytes amongst innate immune cells (fluorescent NP were found mainly in liver F4/80 stained cells (**FigS2D**)), additional staining on myeloid markers were performed (**Fig. 6 and S8**). To start with, it is worth noting that no differences were noticed between mock-infected, PBS and 3TC groups for all following IHC staining, thus suggesting no intrahepatic immune changes for these 3 control groups. In sharp contrast, staining for total macrophage marker F4/80 evidenced an increased number/area of positive immune cells in Pam_3_CSK_4_ treated mice, with highest incidence being observed in NP-Pam_3_CSK_4_ high group; yet differences between all Pam_3_CSK_4_ treated groups were not significative for this marker (**Fig. 6A**). A similar pattern was found for the ubiquitous CD11b marker of monocytes/MDSC (myeloid-derived suppressor cells) with respectively 11-fold vs 13-fold vs 26-fold increase for Pam_3_CSK_4_, NP-Pam_3_CSK_4_ low, and NP-Pam_3_CSK_4_ high groups compared to PBS group (**Fig. 6B**). This time NP-Pam_3_CSK_4_ high group was significantly higher than other Pam_3_CSK_4_ groups, correlating this marker of monocytes (possibly neo-recruited to the parenchyma) to the best antiviral effect seen in the cohort. This high myeloid lineage expression was accompanied by an inverse decline level of liver macrophages (mainly Kupffer cells) within Pam_3_CSK_4_ groups (significative for NP-Pam_3_CSK_4_ high vs free Pam_3_CSK_4_ group, as illustrated by Clec4f staining (**Fig. 6C**). Data from these 3 first panels suggest a strong infiltration of circulating monocytes from the blood to the liver, implying that the TLR1/2 target is well engaged locally. Moreover, MHC II expression was significantly enhanced for NP-Pam_3_CSK_4_ low and NP-Pam_3_CSK_4_ high groups compared to PBS (17-fold) and also free Pam_3_CSK_4_ high (2-fold) groups, indicating an increased M1-like phenotype in NP-Pam_3_CSK_4_ groups (**Fig. 6D**). In an opposite manner, CD206 level, which is indicative of an M2-like phenotype, was decreased in all Pam_3_CSK_4_ groups compared to PBS/3TC groups (**Fig. 6E**). The two last staining suggest that TLR1/2 stimulation favours a pro-inflammatory environment by switching a liver-prone M2-like/tolerogenic phenotype into a M1-like phenotype most likely prompt to viral elimination. Lastly, T-cell markers CD3, CD4 and CD8 were investigated and found increased in all Pam_3_CSK_4_ treated groups, with no significative differences within these groups (**Fig.7 and S9**); this suggest that T-cell responses could also contribute to the overall antiviral observed in all Pam_3_CSK_4_ treated groups, although not fully correlating with subtle variable antiviral effects observed between all Pam_3_CSK_4_ treated groups.

**Figure 6.**
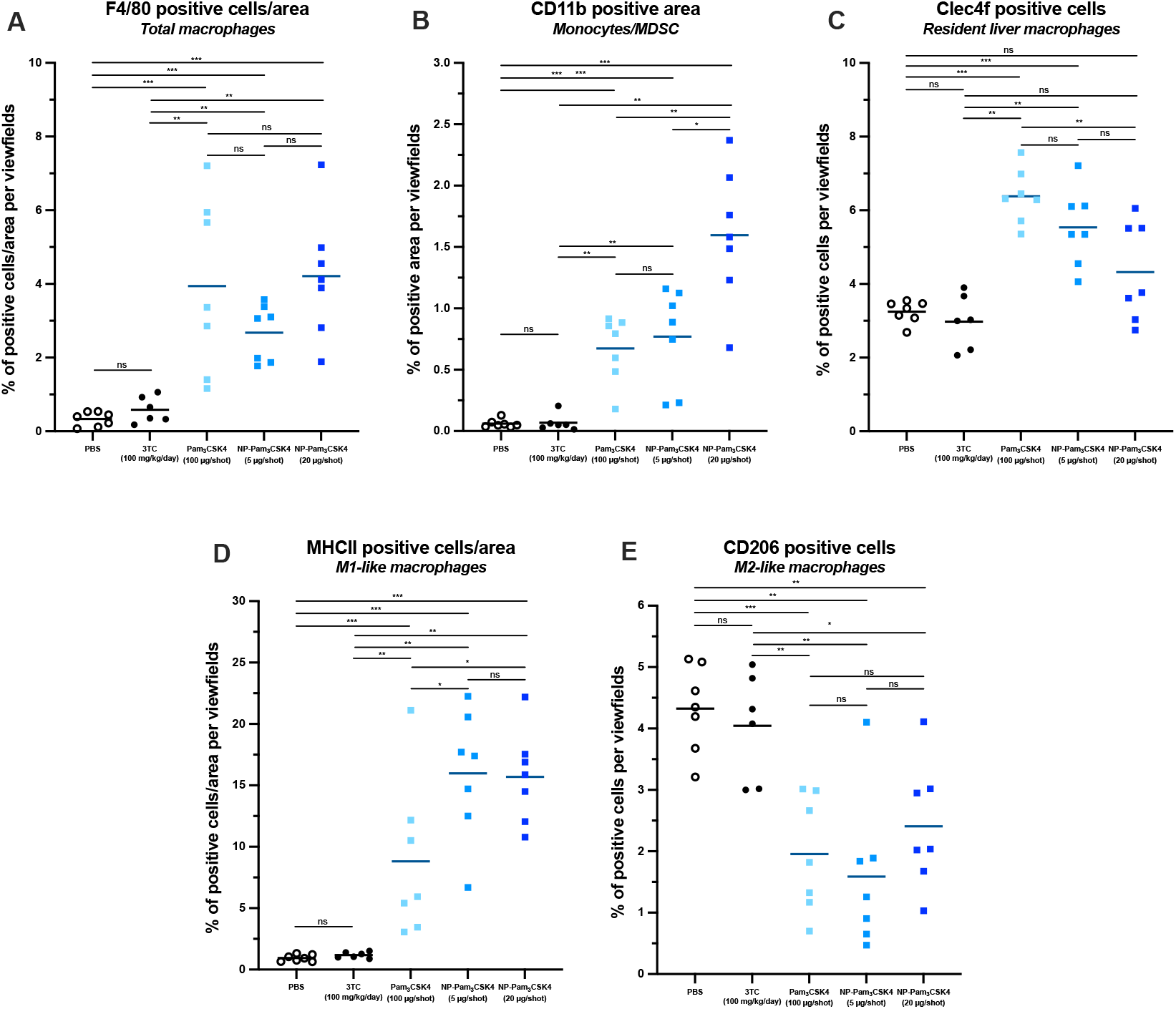
Analysis of myeloid/macrophages markers in liver parenchyma of Pam_3_CSK_4_-treated mice. Immune staining was performed with indicated antibodies on fixed and paraffin-embedded liver slices. Slides were scanned on a SCN400 slide scanner (Leica), and labelled markers were quantified using ImageJ software. (A) Quantification of F4/80 marker. (B) Quantification of CD11b marker. (C) Quantification of Clec4f marker. (D) Quantification of MHCII marker. (E) Quantification of CD206 marker. Threshold for statistical significance was 0.05 for all analysis, with *p < 0.05, **p < 0.01, and ***p < 0.005 ; ns means non-significative.

**Figure 7.**
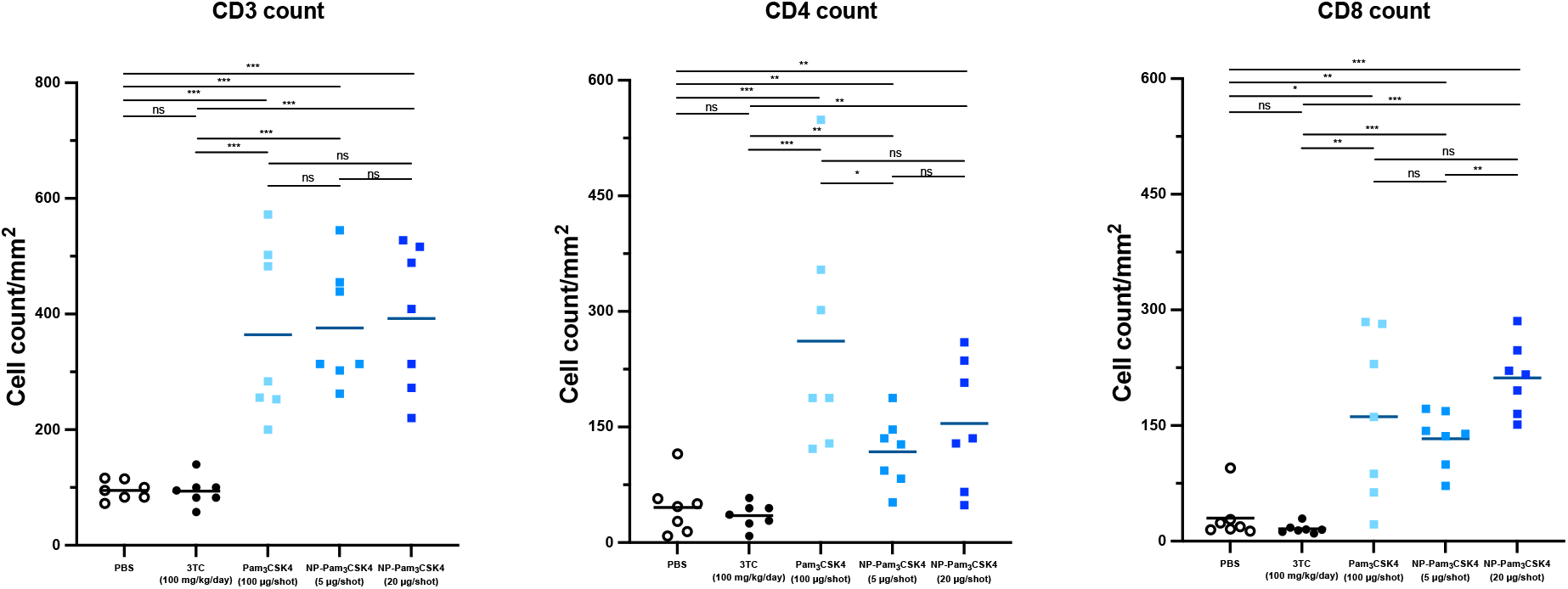
Analysis of T-cell markers in liver parenchyma of Pam_3_CSK_4_-treated mice. Immune staining was performed with indicated antibodies on fixed and paraffin-embedded liver slices. Slides were scanned on a SCN400 slide scanner (Leica), and labelled markers were quantified using ImageJ software. (A) Quantification of CD3 marker. (B) Quantification of CD4 marker. (C) Quantification of CD8 marker. Threshold for statistical significance was 0.05 for all analysis, with *p < 0.05, **p < 0.01, and ***p < 0.005 ; ns means non-significative.

In addition to cell immune staining, targeted RTqPCR to quantify mRNA expression of prototypic proinflammatory genes (IL1β, IL10, TNFα, and IL6) was performed on RNAs extracted from liver pieces (**Fig. 8**). Correlating with results obtained by IHC staining on myeloid inflammatory markers, we found that these genes were upregulated in Pam_3_CSK_4_ groups, compared to PBS and 3TC groups and were overall most expressed in the NP-Pam_3_CSK_4_ high group.

**Figure 8.**
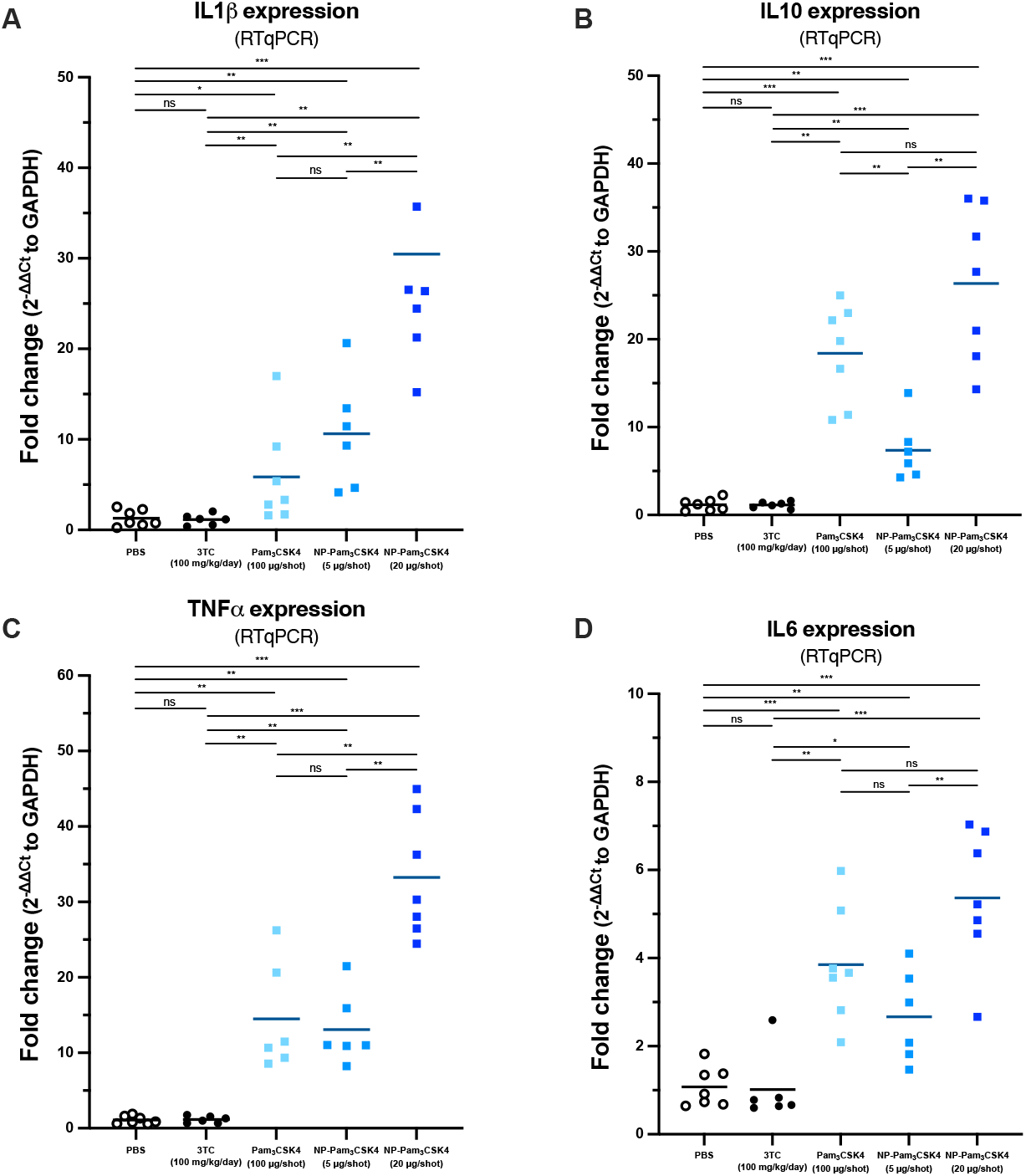
Quantification of prototypic inflammatory genes in liver of Pam_3_CSK_4_-treated mice. RTqPCR were performed with specific primers for indicated genes on RNAs extracted from liver pieces. (A) Quantification of IL1β RNAs. (B) Quantification of IL10 RNAs. (C) Quantification of TNFα RNAs. (D) Quantification of IL6 RNAs. Threshold for statistical significance was 0.05 for all analysis, with *p < 0.05, **p < 0.01, and ***p < 0.005 ; ns means non-significative.

Taken together, TLR1/2 stimulation was found associated with an inflammatory process that include a strong recruitment of inflammatory monocytes and a M1-like polarization/repolarization of these monocytes and resident macrophages, which somehow correlates with the antiviral effect observed. Yet these various immunological patterns did not fully correlate with the antiviral phenotype observed for NP-Pam_3_CSK_4_ groups, confirming that Pam_3_CSK_4_ could also be inducing in a dose-dependent manner a direct antiviral effect within infected hepatocytes, as previously shown *in vitro* (Desmares et al., 2022; Lucifora et al., 2018; Michelet et al., 2022).

## Discussion

To improve the rate of HBV cure, which is rather low with currently approved anti-HBV drugs, it is widely thought that an immune component(s) should be part of a future successful combination strategy. In this respect it is important to keep in mind that the best approved option, to achieve between 3 and 7 % of *functional cure* in real life studies, relies currently on Peg-IFN-a, which is nothing el se than an innate immune component. Yet the systemic administration of Peg-IFN-a, is associated with many side effects that are not well tolerated by patients, who have access to very safe NUCs. In contrast to Peg-IFN-a, NUCs do not lead to a reawakening of immune response against the virus, thus prompting for the R&D alternative innate immune stimulators. Moreover, a specific delivery to the liver of the innate immune component could reduce systemic effect and lead to a safer use.

Several *in vitro* studies had shown that HBV (and HDV) replication and spreading in *bona fide* hepatocytes was more sensitive to “NF-κB inducers” (of both canonical and non-canonical pathways) than to IFN (Isorce et al., 2016; Lucifora et al., 2014; Michelet et al., 2022; Namineni et al., 2020). And amongst TLR agonists capable to engage NF-κB pathways, TLR2-ligands were the most potent to directly inhibit HBV replication in hepatocytes (Desmares et al., 2022; Lucifora et al., 2018), while TLR7 (Niu et al., 2018) and TLR8 (Roca Suarez et al., 2024) agonists were ineffective due to the fact that hepatocytes do not functionally express these two receptors (Faure-Dupuy, Vegna, et al., 2018; Luangsay et al., 2015); TLR7 and TLR8 agonists, such as vesatolimod/GS9620 and selgantolimod/GS9688 mediates their anti-HBV effects by activating initially and respectively pDC and monocytes/macrophages (Amin et al., 2021; Daffis et al., 2021; Mori et al., 2023; Niu et al., 2018; Roca Suarez et al., 2024), which in turn modulates T and B cells activities against HBV *in vivo* (Amin et al., 2021; Daffis et al., 2021; L. Li et al., 2018). Only TLR8 agonists are being pursued in clinical trials against HBV (*Hepatitis B Foundation: Drug Watch*, n.d.). Selgantolimod was recently reported safe in virally-suppressed CHB patients, but demonstrated only a limited efficacy when used alone on top of a NUC (E. J. Gane et al., 2023; H. L. Janssen et al., 2024).

Nanotechnology allows to design new immunomodulatory solutions against hard-to-treat pathogens or tumors by overcoming common challenges encountered with small lipopeptides or molecules, including rapid degradation of drugs, insufficient activation at the site of interest and/or severe side effects in clinical trials (X. Li et al., 2021; Schunke et al., 2023). In this study, aiming at leveraging both direct and indirect antiviral effects for a more potent control of HBV replication *in vivo*, yet mitigating extra liver exposure to the innate immune stimulator, the prototypic TLR1/2 agonist Pam_3_CSK_4_ was vectorized into PLA NP and compared to free form in infected mice. PLA-based technology was chosen based on our previous work and because it is considered very safe. In this study the innocuity and neutrality in the otherwise antiviral phenotypes of PLA-NP were verified. PLA-NP demonstrated an excellent bioavailability and retention rate in the liver, allowing two IV administrations every cycle of 11 days *in vivo*. Besides, the vectorization of Pam_3_CSK_4_ in the case of CHB may sustain the presence of the molecule on-site, therefore enhancing the innate immune activation with a prolonged effect to strengthen the engagement of myeloid and lymphoid cells.

The best antiviral effects were obtained in NP-Pam_3_CSK_4_ groups (5 μg and 20 μg concentrations), with no significative differences between these two NP-Pam_3_CSK_4_ groups at day-90; this indicates that a dose as low as 5 μg/injection (equivalent 200 μg/kg; 8 injections in total over a period of 55 days) of Pam_3_CSK_4_ incorporated into PLA-NP has a far greater effect than a dose of 100 μg/injection of free/non-vectorized Pam_3_CSK_4_. In the first study investigating the effect of various PRR agonists against HBV replication in a transgenic mouse model, Pam_3_CSK_4_ was found ineffective at 200 μg (Isogawa et al., 2005); but it was injected only once and was not protected. In our previous study, a dose escalation administration of free/non-protected Pam_3_CSK_4_ up to 80 μg/injection (*i. e*. 3 cycles of 20, 40 and 80 μg/injection) into HBV-infected liver-humanized mice led only to a mild effect (Lucifora et al., 2018). The vectorisation of Pam_3_CSK_4_ seems to confirm its potential use in future, and targeted delivery plus lowering the concentration of an innate immune stimulator is undeniably desirable to reduce potential safety issues.

NP-Pam_3_CSK_4_ highly decreased viremia of more than 4 log_10_ in maximum at day-90 (end of follow-up) (in two different cohorts). This NP-Pam_3_CSK_4_-driven effect was superior compared to other innate immune stimulators, including i) the TLR7 agonist JNJ-4964 used at 20Ömg/kg once-per-week (*per os*; 500 µg/injection) for 12 weeks (Herschke et al., 2021), ii) an ALPK1 agonist used at 25Öμg/kg once daily for 28 days (*per os*; 625 pg/injection) (Xu et al., 2023), or a STING agonist used at 310 μg/kg once-per-week (*per os*; 7.8 µg/injection) for 10 weeks (Wang et al., 2025).

NP-Pam_3_CSK_4_ moderately decreased HBsAg antigenemia with at best a 0.6 log_10_ reduction (and no anti-HBs seroconversion). Similar weak effect on HBsAg levels were observed with ALPK1 and STING agonists (Wang et al., 2025; Xu et al., 2023), whereas the TLR7 agonist JNJ-4964 led to a more than 2 log_10_ reduction, which was accompanied by anti-HBs seroconversion (Herschke et al., 2021). This limited effect was unexpected because *in vitro* studies had shown a significative effect on HBsAg secretion, as the result of a strong effect on HBV RNA biogenesis (Desmares et al., 2022; Lucifora et al., 2018). Yet these *in vitro* studies highlighted long HBV RNAs (*i. e*. preC and pg RNAs) were particularly impacted by TLR1/2 agonization; as pgRNA is the genomic source for production of rcDNA-containing virions, the *in vivo* preeminent phenotype observed in this study on HBV viremia makes sense and warrant further *in vitro* investigation to work out detailed underlying mechanisms.

Intrahepatic virological analyses aligned rather well with serological ones, revealing increasing antiviral effect from upstream to downstream events in the chain of production of virions (*i. e*. viral episomes --> pgRNA --> t/rcDNA --> secreted DNA; respective panels C, E, D, and A), and a more moderate effect on the chain of production of viral antigens/protein for the 3 Pam_3_CSK_4_ groups; it is worth noting than the trendy orders (in term of antiviral intensity) between 3 Pam_3_CSK_4_ groups were similar for all parameters analysed, thus confirming some dose effect and the quality of analyses. The best antiviral profile was trendy observed in the NP-Pam_3_CSK_4_ high group, although there were no significative differences with NP-Pam_3_CSK_4_ low group.

TLR1/2 receptor engagement in hepatocytes leads to the expression of many NF-κB inducible genes allowing the production of intracellular antiviral effector proteins and secreted cytokines/chemokines. So far, driver effectors of the direct antiviral effect of Pam_3_CSK_4_ on HBV replication in hepatocyte are not known (Desmares et al., 2022); but hepatocyte secreted cytokines (*i. e*. IL6 and TNFα) were shown not contributing to the antiviral effect *in vitro* in the absence of immune cells (Lucifora et al., 2018). TLR1/2 receptor engagement in Kupffer cells/macrophages/myeloid cells leads to the production of various proinflammatory cytokines/chemokines, which are important to coordinate innate and adaptive immune responses. We have shown in this study that PLA-NP are readily taken up by murine liver macrophages upon IV injection; this suggest that NP-Pam_3_CSK_4_ may stimulate liver macrophages and induce proinflammatory cytokine/chemokines liver production. Amongst them IL1β, IL6, and TNFα are known to directly inhibit HBV replication in hepatocytes in the absence of other cells in the *in vitro* culture (Delphin et al., 2021; Faure-Dupuy et al., 2019; Hösel et al., 2009; Isorce et al., 2015, 2016; Watashi et al., 2013; Xia et al., 2016). It is therefore interesting that these 3 cytokines were found in this study expressed in the liver of all Pam_3_CSK_4_ treated mice. Their overall highest expression in the NP-Pam_3_CSK_4_ high group may account at the margin on the trendy better antiviral effect seen in that group of mice. Yet long lasting expression of these genes in the liver could be associated with safety issues, and it is therefore nice to see a decorrelation of their expression with the antiviral effect achieved in NP-Pam_3_CSK_4_ low group.

Pam_3_CSK_4_-induced proinflammatory cytokine/chemokines by liver macrophages was expected to lead to recruitment, liver infiltration and activation of other myeloid cells. Our immune-histochemistry results suggest that it could be indeed the case with increased markers (F4/80 and CD11b) of myeloid/macrophage/monocyte cells quantified in all Pam_3_CSK_4_-treated mice, with a tendency of highest presence in NP-Pam_3_CSK_4_ high group. Interestingly we also found that myeloid/macrophage/monocyte cells detected in all Pam_3_CSK_4_-treated mice were more proinflammatory (M1-like phenotype) and less tolerogenic (M2-like phenotype). This shift in polarisation of liver myeloid/macrophage/monocyte fits well with a possible recovery of immune responses against HBV, knowing that we had previously shown that CHB infection is associated with HBsAg driven M2-like polarisation of Kupffer cells in patients and production of tolerogenic IL10 cytokine (Delphin et al., 2021; Faure-Dupuy, Durantel, et al., 2018; Faure-Dupuy et al., 2019). We also found by IHC staining that T-cell were more abundant in the liver parenchyma of Pam_3_CSK_4_-treated mice, thus suggesting that specific adaptive responses might have occurred and contribute to the overall antiviral effect observed. Yet no significative difference was found between the different Pam_3_CSK_4_ groups, indicating that nano-vectorisation of Pam_3_CSK_4_ did not have an impact. Further investigations on these T cells are yet to be performed to better demonstrate a possible contribution to antiviral effect. Such studies could not be done in the present work and would require running other mouse cohorts. As in our case no HBsAg seroclearance was observed, it is also likely that B cell did not play a major role in the antiviral effect observed; but this should also be better investigated with other mouse cohorts. Other innate immune stimulators (*i. e*. TLR7, ALPK1, and STING agonists), studied in the same AAV-HBV-transduced mouse model, also lead to the recruitment, liver infiltration and activation of myeloid and/or T and/or B cells. However, it is yet difficult to compare all these studies and get a clear understanding on the drivers (direct or indirect) at work to explain variable antiviral activity with these different innate immune stimulators. Various agonists should be investigated side by side, in the same model and with the same reads-out for a better comparison of their direct and immune-driven antiviral and safety profiles.

Both myeloid/macrophage/monocyte and T-cells IHC staining have shown a clusterization of these immune cells in the parenchyma of Pam_3_CSK_4_-treated mice; these “clusters” resembled previously characterized intrahepatic myeloid cell aggregates for T cell expansion (iMATE), that were formed after TLR9 and TLR4 immunotherapy (Huang et al., 2013) or tertiary lymphoid structures (TLS) (Guillaume et al., 2025). These structures enable expansion of liver cytotoxic T cell (CTL) population in close contact with myeloid/macrophage/monocytes cells. It is tempting to speculate that TLR1/2 agonists could induce this kind of structures, which could in turn correlates with antiviral effect observed. But this was beyond the scope of the present study and will require further investigation. Further immune assays, as well as spatial transcriptomics/proteomics would be needed to investigate the exact nature of these Pam_3_CSK_4_-induced immune clusters as well as the specificity and activation profile of the lymphoid compartment.

To conclude, we have demonstrated in this work that the Pam_3_CSK_4_ agonist can induce a potent viro-suppression (*i. e*. strong decrease in viremia) when used in monotherapy in AAV-HBV infected C57BL6 mice and that its vectorisation into PLA NP is safe and allow to reduce the dose needed for antiviral effect to as low as 200 μg/kg per injection. Intrahepatic antiviral effect was aligned with that measured in the serum. NP-Pam_3_CSK_4_ low treatment was as antiviral as NP-Pam_3_CSK_4_ high one and is anticipated to be a safer option. Yet both NP-Pam_3_CSK_4_ groups were associated with similar immune profiles, characterized by i) myeloid/macrophage/monocytes (and T-cell) cells infiltration and activation in liver parenchyma, ii) an overall inflammation prone to antiviral activity, in link with a repolarisation of myeloid/macrophage/monocytes from tolerogenic to inflammatory phenotype and expression of proinflammatory cytokines, iii) a compensated liver turnover, ending in a slight hepatomegaly at euthanasia, but similar weight of mice. This vectorized TLR1/2 agonist should be investigated further to determine whether it could be used in combination therapy and/or as an adjuvant for boosting immune therapeutic vaccine. And more toxicologic analyses are warranted to firmly demonstrate whether it could be moved forward to human use.

## Abbreviations

AAV: adeno-associated virus
cccDNA: covalently closed circular DNA
CHB: chronic hepatitis B virus infection
CLIA: chemiluminescence immunoassay
CTL: cytotoxic T cells
HBeAg: HBV antigen e
HBsAg: HBV surface antigen
HBV: hepatitis B virus
H&E: hematoxylin and eosin
IV: intravenous
IFN: interferon
IHC: immunohistochemistry
iMATEs: intrahepatic myeloid cell aggregates for T cell expansion
KO: knocked-out
LLOQ: lower limit of quantification
MHCII: major histocompatibility complex type II molecules
NF-κB: nuclear factor kappa-light-chain-enhancer of activated B cells
NK cells: natural killer cells
NP: nanoparticle
NP high: NP-Pam3CSK4 high dose (20 μg/injection)
NP low: NP-Pam3CSK4 low dose (5 μg/injection)
NUCs: nucleos(t)ide analogues
Peg-IFN-a: pegylated IFN alfa
pgRNA: HBV pregenomic RNA
PLA: polylactic acid
SPF: specific pathogen-free
SVP: HBV subviral particles
TCR: T cell receptor
TLR: Toll-Like-Receptor
versus: vs
vge: virus genome equivalent
wT: wild type.

## Acknowledgements

The authors would like to thank staff from animal facilities for their technical help in mouse handling. We acknowledge the contributions of the CELPHEDIA Infrastructure (http://www.celphedia.eu/), especially the center AniRA in Lyon. This work was supported by two grants from the CSS12 of ANRS-MIE (Agence Nationale pour la Recherche sur le Sida, les Hépatites Vials & les Maladies Infectieuses Émergentes, ECTZ65108 and ECTZ137751), an Infect-Era European grant operated by ANR (Agence Nationale pour la Recherche; ANR 16-IFEC-0005-01), MSD-Avenir France (Hit-Hidden HBV program), as well as core financial support from INSERM, CNRS, and UCBL1. Bernard Verrier has founded Adjuvatis SA. All other authors have no conflict of interest to declare.

## References

Amin, O. E., Colbeck, E. J., Daffis, S., Khan, S., Ramakrishnan, D., Pattabiraman, D., Chu, R., Micolochick Steuer, H., Lehar, S., Peiser, L., Palazzo, A., Frey, C., Davies, J., Javanbakht, H., Rosenberg, W. M. C., Fletcher, S. P., Maini, M. K., & Pallett, L. J. (2021). Therapeutic Potential of TLR8 Agonist GS-9688 (Selgantolimod) in Chronic Hepatitis B: Remodeling of Antiviral and Regulatory Mediators. Hepatology, 74(1), 55–71. 10.1002/hep.31695

Bankhead, P., Loughrey, M. B., Fernández, J. A., Dombrowski, Y., McArt, D. G., Dunne, P. D., McQuaid, S., Gray, R. T., Murray, L. J., Coleman, H. G., James, J. A., Salto-Tellez, M., & Hamilton, P. W. (2017). QuPath: Open source software for digital pathology image analysis. Scientific Reports, 7(1), 16878. 10.1038/s41598-017-17204-5

Boni, C., Vecchi, A., Rossi, M., Laccabue, D., Giuberti, T., Alfieri, A., Lampertico, P., Grossi, G., Facchetti, F., Brunetto, M. R., Coco, B., Cavallone, D., Mangia, A., Santoro, R., Piazzolla, V., Lau, A., Gaggar, A., Subramanian, G. M., & Ferrari, C. (2018). TLR7 Agonist Increases Responses of Hepatitis B Virus-Specific T Cells and Natural Killer Cells in Patients With Chronic Hepatitis B Treated With Nucleos(T)Ide Analogues. Gastroenterology, 154(6), 1764-1777.e7. 10.1053/j.gastro.2018.01.030

Bouquet, J., Speer, S. D., Koenig, A., Livingston, C. M., Savarese, M., Lau, C., Angelini, E., Galwey, N., Yates, P., Maynard, L., You, S., Cremer, J., Paff, M., Theodore, D., Elston, R., & Walker, J. (2026). Single nucleotide polymorphisms in the bepirovirsen binding site have limited impact on treatment response in chronic hepatitis B. Journal of Hepatology, 84(5), 887–897. 10.1016/j.jhep.2025.12.025

Daffis, S., Balsitis, S., Chamberlain, J., Zheng, J., Santos, R., Rowe, W., Ramakrishnan, D., Pattabiraman, D., Spurlock, S., Chu, R., Kang, D., Mish, M., Ramirez, R., Li, L., Li, B., Ma, S., Hung, M., Voitenleitner, C., Yon, C., … Fletcher, S. P. (2021). Toll-Like Receptor 8 Agonist GS-9688 Induces Sustained Efficacy in the Woodchuck Model of Chronic Hepatitis B. Hepatology, 73(1), 53–67. 10.1002/hep.31255

Delphin, M., Faure-Dupuy, S., Isorce, N., Rivoire, M., Salvetti, A., Durantel, D., & Lucifora, J. (2021). Inhibitory Effect of IL-1β on HBV and HDV Replication and HBs Antigen-Dependent Modulation of Its Secretion by Macrophages. Viruses, 14(1), 65. 10.3390/v14010065

Desmares, M., Delphin, M., Chardès, B., Pons, C., Riedinger, J., Michelet, M., Rivoire, M., Verrier, B., Salvetti, A., Lucifora, J., & Durantel, D. (2022). Insights on the antiviral mechanisms of action of the TLR1/2 agonist Pam3CSK4 in hepatitis B virus (HBV)-infected hepatocytes. Antiviral Research, 206, 105386. 10.1016/j.antiviral.2022.105386

Dion, S., Bourgine, M., Godon, O., Levillayer, F., & Michel, M.-L. (2013). Adeno-associated virus-mediated gene transfer leads to persistent hepatitis B virus replication in mice expressing HLA-A2 and HLA-DR1 molecules. Journal of Virology, 87(10), 5554–5563. 10.1128/JVI.03134-12

Dou, Y., Jansen, D. T. S. L., van den Bosch, A., de Man, R. A., van Montfoort, N., Araman, C., van Kasteren, S. I., Zom, G. G., Krebber, W.-J., Melief, C. J. M., Woltman, A. M., & Buschow, S. I. (2020). Design of TLR2-ligand-synthetic long peptide conjugates for therapeutic vaccination of chronic HBV patients. Antiviral Research, 178, 104746. 10.1016/j.antiviral.2020.104746

Dusheiko, G., Agarwal, K., & Maini, M. K. (2023). New Approaches to Chronic Hepatitis B. The New England Journal of Medicine, 388(1), 55–69. 10.1056/NEJMra2211764

European Association for the Study of the Liver. (2025). EASL Clinical Practice Guidelines on the management of hepatitis B virus infection. Journal of Hepatology, 83(2), 502–583. 10.1016/j.jhep.2025.03.018

Faure-Dupuy, S., Delphin, M., Aillot, L., Dimier, L., Lebossé, F., Fresquet, J., Parent, R., Matter, M. S., Rivoire, M., Bendriss-Vermare, N., Salvetti, A., Heide, D., Flores, L., Klumpp, K., Lam, A., Zoulim, F., Heikenwälder, M., Durantel, D., & Lucifora, J. (2019). Hepatitis B virus-induced modulation of liver macrophage function promotes hepatocyte infection. Journal of Hepatology, 71(6), 1086–1098. 10.1016/j.jhep.2019.06.032

Faure-Dupuy, S., Durantel, D., & Lucifora, J. (2018). Liver macrophages: Friend or foe during hepatitis B infection? Liver International: Official Journal of the International Association for the Study of the Liver, 38(10), 1718–1729. 10.1111/liv.13884

Faure-Dupuy, S., Vegna, S., Aillot, L., Dimier, L., Esser, K., Broxtermann, M., Bonnin, M., Bendriss-Vermare, N., Rivoire, M., Passot, G., Lesurtel, M., Mabrut, J.-Y., Ducerf, C., Salvetti, A., Protzer, U., Zoulim, F., Durantel, D., & Lucifora, J. (2018). Characterization of Pattern Recognition Receptor Expression and Functionality in Liver Primary Cells and Derived Cell Lines. Journal of Innate Immunity, 10(4), 339–348. 10.1159/000489966

Frederick, J. M. (1987). The emergence of GABA-accumulating neurons during retinal histogenesis in the embryonic chick. Experimental Eye Research, 45(6), 933–945. 10.1016/s0014-4835(87)80107-0

Gane, E. J., Dunbar, P. R., Brooks, A. E., Zhang, F., Chen, D., Wallin, J. J., van Buuren, N., Arora, P., Fletcher, S. P., Tan, S. K., Yang, J. C., Gaggar, A., Kottilil, S., & Tang, L. (2023). Safety and efficacy of the oral TLR8 agonist selgantolimod in individuals with chronic hepatitis B under viral suppression. Journal of Hepatology, 78(3), 513–523. 10.1016/j.jhep.2022.09.027

Gane, E., Pastagia, M., Schwertschlag, U., De Creus, A., Schwabe, C., Vandenbossche, J., Slaets, L., Fevery, B., Smyej, I., Wu, L. S., Li, R., Siddiqui, S., Oey, A., Musto, C., & Van Remoortere, P. (2021). Safety, tolerability, pharmacokinetics, and pharmacodynamics of oral JNJ-64794964, a TLR-7 agonist, in healthy adults. Antiviral Therapy, 26(3–5), 58–68. 10.1177/13596535211056581

Gehring, A. J., & Protzer, U. (2019). Targeting Innate and Adaptive Immune Responses to Cure Chronic HBV Infection. Gastroenterology, 156(2), 325–337. 10.1053/j.gastro.2018.10.032

Guillaume, S. M., Beccaria, C. G., Iannacone, M., & Linterman, M. A. (2025). Tertiary Lymphoid Structures Across Organs: Context, Composition, and Clinical Levers. Immunological Reviews, 335(1), e70063. 10.1111/imr.70063 Hepatitis B Foundation: Drug Watch. (n.d.). Retrieved April 23, 2026, from https://www.hepb.org/treatment-and-management/drug-watch/

Herschke, F., Li, C., Zhu, R., Han, Q., Wu, Q., Lu, Q., Barale-Thomas, E., De Jonghe, S., Lin, T.-I., & De Creus, A. (2021). JNJ-64794964 (AL-034/TQ-A3334), a TLR7 agonist, induces sustained anti-HBV activity in AAV/HBV mice via non-cytolytic mechanisms. Antiviral Research, 196, 105196. 10.1016/j.antiviral.2021.105196

Hong, J., & Rajwanshi, V. K. (2025). Antisense oligonucleotides as drugs with both direct and indirect antiviral actions. Antiviral Research, 240, 106219. 10.1016/j.antiviral.2025.106219

Hösel, M., Quasdorff, M., Wiegmann, K., Webb, D., Zedler, U., Broxtermann, M., Tedjokusumo, R., Esser, K., Arzberger, S., Kirschning, C. J., Langenkamp, A., Falk, C., Büning, H., Rose-John, S., & Protzer, U. (2009). Not interferon, but interleukin-6 controls early gene expression in hepatitis B virus infection. Hepatology, 50(6), 1773–1782. 10.1002/hep.23226

Hu, Y., Zhang, H., Wu, M., Liu, J., Li, X., Zhu, X., Li, C., Chen, H., Liu, C., Niu, J., & Ding, Y. (2021). Safety, pharmacokinetics and pharmacodynamics of TQ-A3334, an oral toll-like receptor 7 agonist in healthy individuals. Expert Opinion on Investigational Drugs, 30(3), 263–269. 10.1080/13543784.2021.1873275

Huang, L.-R., Wohlleber, D., Reisinger, F., Jenne, C. N., Cheng, R.-L., Abdullah, Z., Schildberg, F. A., Odenthal, M., Dienes, H.-P., van Rooijen, N., Schmitt, E., Garbi, N., Croft, M., Kurts, C., Kubes, P., Protzer, U., Heikenwalder, M., & Knolle, P. A. (2013). Intrahepatic myeloid-cell aggregates enable local proliferation of CD8(+) T cells and successful immunotherapy against chronic viral liver infection. Nature Immunology, 14(6), 574–583. 10.1038/ni.2573

Isogawa, M., Robek, M. D., Furuichi, Y., & Chisari, F. V. (2005). Toll-like receptor signaling inhibits hepatitis B virus replication in vivo. Journal of Virology, 79(11), 7269–7272. 10.1128/JVI.79.11.7269-7272.2005

Isorce, N., Lucifora, J., Zoulim, F., & Durantel, D. (2015). Immune-modulators to combat hepatitis B virus infection: From IFN-α to novel investigational immunotherapeutic strategies. Antiviral Research, 122, 69–81. 10.1016/j.antiviral.2015.08.008

Isorce, N., Testoni, B., Locatelli, M., Fresquet, J., Rivoire, M., Luangsay, S., Zoulim, F., & Durantel, D. (2016). Antiviral activity of various interferons and pro-inflammatory cytokines in non-transformed cultured hepatocytes infected with hepatitis B virus. Antiviral Research, 130, 36–45. 10.1016/j.antiviral.2016.03.008

Janssen, H. L. A., Brunetto, M. R., Kim, Y. J., Ferrari, C., Massetto, B., Nguyen, A.-H., Joshi, A., Woo, J., Lau, A. H., Gaggar, A., Subramanian, G. M., Yoshida, E. M., Ahn, S. H., Tsai, N. C. S., Fung, S., & Gane, E. J. (2018). Safety, efficacy and pharmacodynamics of vesatolimod (GS-9620) in virally suppressed patients with chronic hepatitis B. Journal of Hepatology, 68(3), 431–440. 10.1016/j.jhep.2017.10.027

Janssen, H. L., Lim, Y.-S., Kim, H. J., Sowah, L., Tseng, C.-H., Coffin, C. S., Elkhashab, M., Ahn, S. H., Nguyen, A.-H., Chen, D., Wallin, J. J., Fletcher, S. P., McDonald, C., Yang, J. C., Gaggar, A., Brainard, D. M., Fung, S., Kim, Y. J., Kao, J.-H., … Dunbar, P. R. (2024). Safety, pharmacodynamics, and antiviral activity of selgantolimod in viremic patients with chronic hepatitis B virus infection. JHEP Reports: Innovation in Hepatology, 6(2), 100975. 10.1016/j.jhepr.2023.100975

Ko, C., Su, J., Festag, J., Bester, R., Kosinska, A. D., & Protzer, U. (2021). Intramolecular recombination enables the formation of hepatitis B virus (HBV) cccDNA in mice after HBV genome transfer using recombinant AAV vectors. Antiviral Research, 194, 105140. 10.1016/j.antiviral.2021.105140

Kosinska, A. D., Moeed, A., Kallin, N., Festag, J., Su, J., Steiger, K., Michel, M.-L., Protzer, U., & Knolle, P. A. (2019). Synergy of therapeutic heterologous prime-boost hepatitis B vaccination with CpG-application to improve immune control of persistent HBV infection. Scientific Reports, 9(1), 10808. 10.1038/s41598-019-47149-w

Lamalle-Bernard, D., Munier, S., Compagnon, C., Charles, M.-H., Kalyanaraman, V. S., Delair, T., Verrier, B., & Ataman-Onal, Y. (2006). Coadsorption of HIV-1 p24 and gp120 proteins to surfactant-free anionic PLA nanoparticles preserves antigenicity and immunogenicity. Journal of Controlled Release: Official Journal of the Controlled Release Society, 115(1), 57–67. 10.1016/j.jconrel.2006.07.006

Lamrayah, M., Charriaud, F., Desmares, M., Coiffier, C., Megy, S., Colomb, E., Terreux, R., Lucifora, J., Durantel, D., & Verrier, B. (2023). Induction of a strong and long-lasting neutralizing immune response by dPreS1-TLR2 agonist nanovaccine against hepatitis B virus. Antiviral Research, 209, 105483. 10.1016/j.antiviral.2022.105483

Lamrayah, M., Charriaud, F., Hu, S., Megy, S., Terreux, R., & Verrier, B. (2019). Molecular modelling of TLR agonist Pam3CSK4 entrapment in PLA nanoparticles as a tool to explain loading efficiency and functionality. International Journal of Pharmaceutics, 568, 118569. 10.1016/j.ijpharm.2019.118569

Li, L., Barry, V., Daffis, S., Niu, C., Huntzicker, E., French, D. M., Mikaelian, I., Lanford, R. E., Delaney, W. E., & Fletcher, S. P. (2018). Anti-HBV response to toll-like receptor 7 agonist GS-9620 is associated with intrahepatic aggregates of T cells and B cells. Journal of Hepatology, 68(5), 912–921. 10.1016/j.jhep.2017.12.008

Li, X., Liu, S., Yin, P., & Chen, K. (2021). Enhanced Immune Responses by Virus-Mimetic Polymeric Nanostructures Against Infectious Diseases. Frontiers in Immunology, 12, 804416. 10.3389/fimmu.2021.804416

Luangsay, S., Ait-Goughoulte, M., Michelet, M., Floriot, O., Bonnin, M., Gruffaz, M., Rivoire, M., Fletcher, S., Javanbakht, H., Lucifora, J., Zoulim, F., & Durantel, D. (2015). Expression and functionality of Toll- and RIG-like receptors in HepaRG cells. Journal of Hepatology, 63(5), 1077–1085. 10.1016/j.jhep.2015.06.022

Lucifora, J., Bonnin, M., Aillot, L., Fusil, F., Maadadi, S., Dimier, L., Michelet, M., Floriot, O., Ollivier, A., Rivoire, M., Ait-Goughoulte, M., Daffis, S., Fletcher, S. P., Salvetti, A., Cosset, F.-L., Zoulim, F., & Durantel, D. (2018). Direct antiviral properties of TLR ligands against HBV replication in immune-competent hepatocytes. Scientific Reports, 8(1), 5390. 10.1038/s41598-018-23525-w

Lucifora, J., Salvetti, A., Marniquet, X., Mailly, L., Testoni, B., Fusil, F., Inchauspé, A., Michelet, M., Michel, M.-L., Levrero, M., Cortez, P., Baumert, T. F., Cosset, F.-L., Challier, C., Zoulim, F., & Durantel, D. (2017). Detection of the hepatitis B virus (HBV) covalently-closed-circular DNA (cccDNA) in mice transduced with a recombinant AAV-HBV vector. Antiviral Research, 145, 14–19. 10.1016/j.antiviral.2017.07.006

Lucifora, J., Xia, Y., Reisinger, F., Zhang, K., Stadler, D., Cheng, X., Sprinzl, M. F., Koppensteiner, H., Makowska, Z., Volz, T., Remouchamps, C., Chou, W.-M., Thasler, W. E., Hüser, N., Durantel, D., Liang, T. J., Münk, C., Heim, M. H., Browning, J. L., … Protzer, U. (2014). Specific and nonhepatotoxic degradation of nuclear hepatitis B virus cccDNA. Science, 343(6176), 1221–1228. 10.1126/science.1243462

Ma, Z., Liu, J., Wu, W., Zhang, E., Zhang, X., Li, Q., Zelinskyy, G., Buer, J., Dittmer, U., Kirschning, C. J., & Lu, M. (2017). The IL-1R/TLR signaling pathway is essential for efficient CD8+ T-cell responses against hepatitis B virus in the hydrodynamic injection mouse model. Cellular & Molecular Immunology, 14(12), 997–1008. 10.1038/cmi.2017.43

Martin, P., Dubois, C., Jacquier, E., Dion, S., Mancini-Bourgine, M., Godon, O., Kratzer, R., Lelu-Santolaria, K., Evlachev, A., Meritet, J.-F., Schlesinger, Y., Villeval, D., Strub, J.-M., Van Dorsselaer, A., Marchand, J.-B., Geist, M., Brandely, R., Findeli, A., Boukhebza, H., … Inchauspé, G. (2015). TG1050, an immunotherapeutic to treat chronic hepatitis B, induces robust T cells and exerts an antiviral effect in HBV-persistent mice. Gut, 64(12), 1961–1971. 10.1136/gutjnl-2014-308041

Michelet, M., Alfaiate, D., Chardès, B., Pons, C., Faure-Dupuy, S., Engleitner, T., Farhat, R., Riedl, T., Legrand, A.-F., Rad, R., Rivoire, M., Zoulim, F., Heikenwälder, M., Salvetti, A., Durantel, D., & Lucifora, J. (2022). Inducers of the NF-κB pathways impair hepatitis delta virus replication and strongly decrease progeny infectivity in vitro. JHEP Reports: Innovation in Hepatology, 4(3), 100415. 10.1016/j.jhepr.2021.100415

Mori, T., Yoshio, S., Yoshikawa, S., Tsustui, Y., Sakata, T., Yoshida, Y., Sakamoto, Y., Kawai, H., Osawa, Y., Yamazoe, T., Aoki, Y., Fletcher, S. P., & Kanto, T. (2023). Toll-like receptor 7 agonist, GS-986, is an immune-stimulant inducing follicular helper T cells and expanding HBs antigen-specific B cells in vitro. Liver International: Official Journal of the International Association for the Study of the Liver, 43(6), 1213–1224. 10.1111/liv.15568

Naghib, M., Kariminik, A., & Kazemi Arababadi, M. (2022). TLR2, as a Pathogen Recognition Receptor, Plays Critical Roles in Hepatitis B Outcome. Viral Immunology, 35(1), 15–23. 10.1089/vim.2021.0141

Namineni, S., O’Connor, T., Faure-Dupuy, S., Johansen, P., Riedl, T., Liu, K., Xu, H., Singh, I., Shinde, P., Li, F., Pandyra, A., Sharma, P., Ringelhan, M., Muschaweckh, A., Borst, K., Blank, P., Lampl, S., Neuhaus, K., Durantel, D., … Heikenwalder, M. (2020). A dual role for hepatocyte-intrinsic canonical NF-κB signaling in virus control. Journal of Hepatology, 72(5), 960–975. 10.1016/j.jhep.2019.12.019

Niu, C., Li, L., Daffis, S., Lucifora, J., Bonnin, M., Maadadi, S., Salas, E., Chu, R., Ramos, H., Livingston, C. M., Beran, R. K., Garg, A. V., Balsitis, S., Durantel, D., Zoulim, F., Delaney, W. E., & Fletcher, S. P. (2018). Toll-like receptor 7 agonist GS-9620 induces prolonged inhibition of HBV via a type I interferon-dependent mechanism. Journal of Hepatology, 68(5), 922–931. 10.1016/j.jhep.2017.12.007

Peres, C., Matos, A. I., Conniot, J., Sainz, V., Zupančič, E., Silva, J. M., Graça, L., Sá Gaspar, R., Préat, V., & Florindo, H. F. (2017). Poly(lactic acid)-based particulate systems are promising tools for immune modulation. Acta Biomaterialia, 48, 41–57. 10.1016/j.actbio.2016.11.012

Roca Suarez, A. A., Plissonnier, M.-L., Grand, X., Michelet, M., Giraud, G., Saez-Palma, M., Dubois, A., Heintz, S., Diederichs, A., Van Renne, N., Vanwolleghem, T., Daffis, S., Li, L., Kolhatkar, N., Hsu, Y.-C., Wallin, J. J., Lau, A. H., Fletcher, S. P., Rivoire, M., … Zoulim, F. (2024). TLR8 agonist selgantolimod regulates Kupffer cell differentiation status and impairs HBV entry into hepatocytes via an IL-6-dependent mechanism. Gut, 73(12), 2012–2022. 10.1136/gutjnl-2023-331396

Sacherl, J., Kosinska, A. D., Kemter, K., Kächele, M., Laumen, S. C., Kerth, H. A., Öz, E. A., Wolff, L. S., Su, J., Essbauer, S., Sutter, G., Scholz, M., Singethan, K., Altrichter, J., & Protzer, U. (2023). Efficient stabilization of therapeutic hepatitis B vaccine components by amino-acid formulation maintains its potential to break immune tolerance. JHEP Reports: Innovation in Hepatology, 5(2), 100603. 10.1016/j.jhepr.2022.100603

Schmidt, U., Weigert, M., Broaddus, C., & Myers, G. (2018). Cell Detection with Star-convex Polygons (Vol. 11071, pp. 265–273). 10.1007/978-3-030-00934-2_30

Schunke, J., Mailänder, V., Landfester, K., & Fichter, M. (2023). Delivery of Immunostimulatory Cargos in Nanocarriers Enhances Anti-Tumoral Nanovaccine Efficacy. International Journal of Molecular Sciences, 24(15), 12174. 10.3390/ijms241512174

Shakya, A. K., Al-Sulaibi, M., Naik, R. R., Nsairat, H., Suboh, S., & Abulaila, A. (2023). Review on PLGA Polymer Based Nanoparticles with Antimicrobial Properties and Their Application in Various Medical Conditions or Infections. Polymers, 15(17), 3597. 10.3390/polym15173597

Wang, Y., Guo, L., Shi, J., Li, J., Wen, Y., Gu, G., Cui, J., Feng, C., Jiang, M., Fan, Q., Tang, J., Chen, S., Zhang, J., Zheng, X., Pan, M., Li, X., Sun, Y., Zhang, Z., Li, X., … Li, F. (2023). Interferon stimulated immune profile changes in a humanized mouse model of HBV infection. Nature Communications, 14(1), 7393. 10.1038/s41467-023-43078-5

Wang, Y., Wu, S., Li, X., Qiao, L., Wang, H., Yang, G., Yan, H., Wang, K., Jiang, J.-D., & Li, Y. (2025). Antiviral and immune modulatory activities of STING agonists in a mouse model of persistent hepatitis B virus infection. PLoS Pathogens, 21(12), e1013709. 10.1371/journal.ppat.1013709

Watashi, K., Liang, G., Iwamoto, M., Marusawa, H., Uchida, N., Daito, T., Kitamura, K., Muramatsu, M., Ohashi, H., Kiyohara, T., Suzuki, R., Li, J., Tong, S., Tanaka, Y., Murata, K., Aizaki, H., & Wakita, T. (2013). Interleukin-1 and tumor necrosis factor-α trigger restriction of hepatitis B virus infection via a cytidine deaminase activation-induced cytidine deaminase (AID). The Journal of Biological Chemistry, 288(44), 31715–31727. 10.1074/jbc.M113.501122

Wolf, M. J., Adili, A., Piotrowitz, K., Abdullah, Z., Boege, Y., Stemmer, K., Ringelhan, M., Simonavicius, N., Egger, M., Wohlleber, D., Lorentzen, A., Einer, C., Schulz, S., Clavel, T., Protzer, U., Thiele, C., Zischka, H., Moch, H., Tschöp, M., … Heikenwalder, M. (2014). Metabolic activation of intrahepatic CD8+ T cells and NKT cells causes nonalcoholic steatohepatitis and liver cancer via cross-talk with hepatocytes. Cancer Cell, 26(4), 549–564. 10.1016/j.ccell.2014.09.003

Xia, Y., Stadler, D., Lucifora, J., Reisinger, F., Webb, D., Hösel, M., Michler, T., Wisskirchen, K., Cheng, X., Zhang, K., Chou, W.-M., Wettengel, J. M., Malo, A., Bohne, F., Hoffmann, D., Eyer, F., Thimme, R., Falk, C. S., Thasler, W. E., … Protzer, U. (2016). Interferon-γ and Tumor Necrosis Factor-α Produced by T Cells Reduce the HBV Persistence Form, cccDNA, Without Cytolysis. Gastroenterology, 150(1), 194–205. 10.1053/j.gastro.2015.09.026

Xu, C., Fan, J., Liu, D., Tuerdi, A., Chen, J., Wei, Y., Pan, Y., Dang, H., Wei, X., Yousif, A. S., Yogaratnam, J., Zhou, Q., Lichenstein, H., & Xu, T. (2023). Alpha-kinase 1 (ALPK1) agonist DF-006 demonstrates potent efficacy in mouse and primary human hepatocyte (PHH) models of hepatitis B. Hepatology, 77(1), 275–289. 10.1002/hep.32614

Yuen, M.-F., Balabanska, R., Cottreel, E., Chen, E., Duan, D., Jiang, Q., Patil, A., Triyatni, M., Upmanyu, R., Zhu, Y., Canducci, F., & Gane, E. J. (2023). TLR7 agonist RO7020531 versus placebo in healthy volunteers and patients with chronic hepatitis B virus infection: A randomised, observer-blind, placebo-controlled, phase 1 trial. The Lancet. Infectious Diseases, 23(4), 496–507. 10.1016/S1473-3099(22)00727-7

Zhang, T. P., & Terrault, N. A. (2024). Peginterferon as Part of a Functional Cure Strategy for Hepatitis B: Is the Juice Worth the Squeeze? Journal of Clinical and Experimental Hepatology, 14(1), 101300. 10.1016/j.jceh.2023.10.092

Zhao, Q., Liu, H., Tang, L., Wang, F., Tolufashe, G., Chang, J., & Guo, J.-T. (2024). Mechanism of interferon alpha therapy for chronic hepatitis B and potential approaches to improve its therapeutic efficacy. Antiviral Research, 221, 105782. 10.1016/j.antiviral.2023.105782

