## Supplementary figures and images for "Preclinical antiviral study of a liver-targeted TLR1/2 agonist in an immune-competent mouse model of HBV infection"

### Supplementary Figures

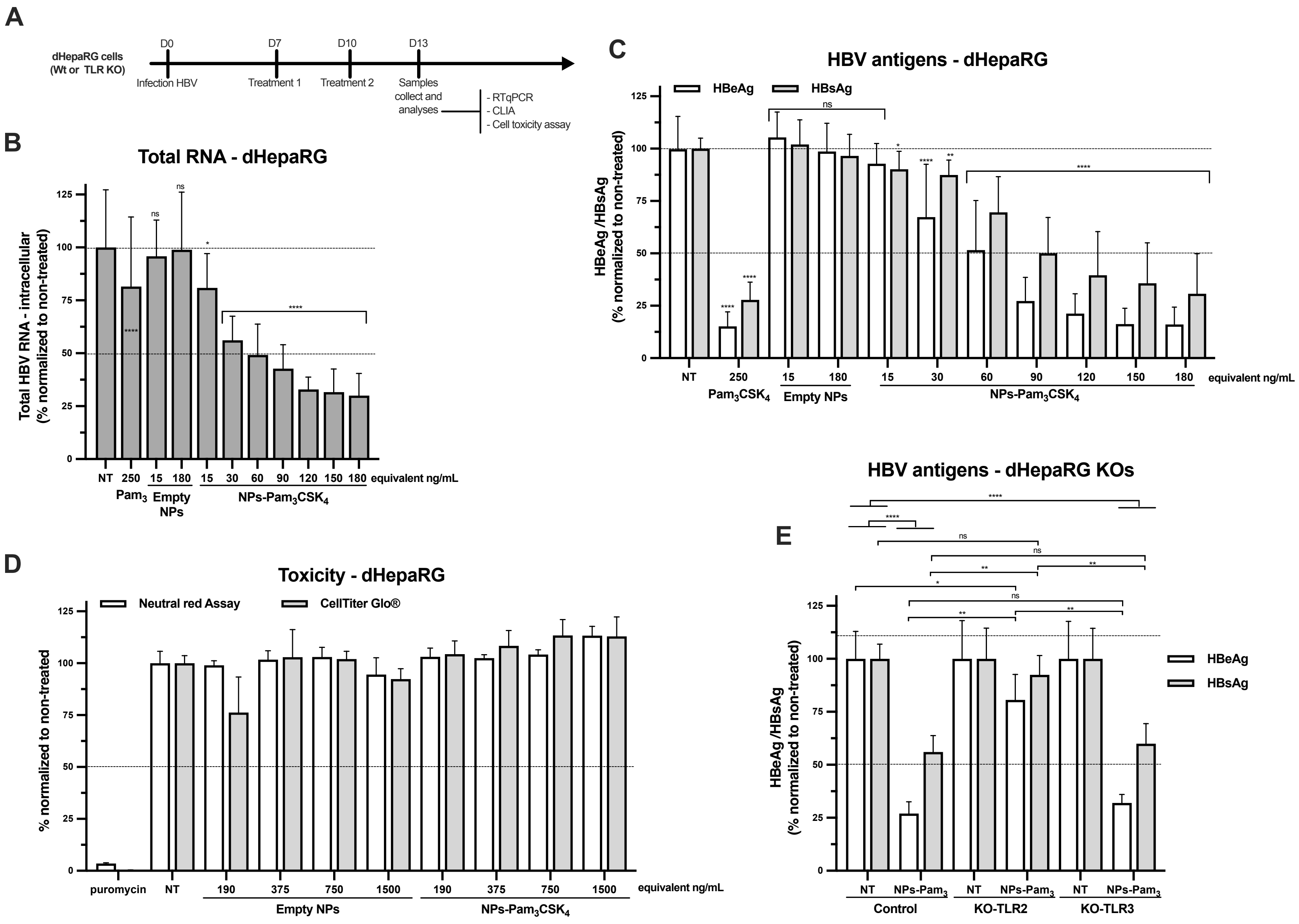

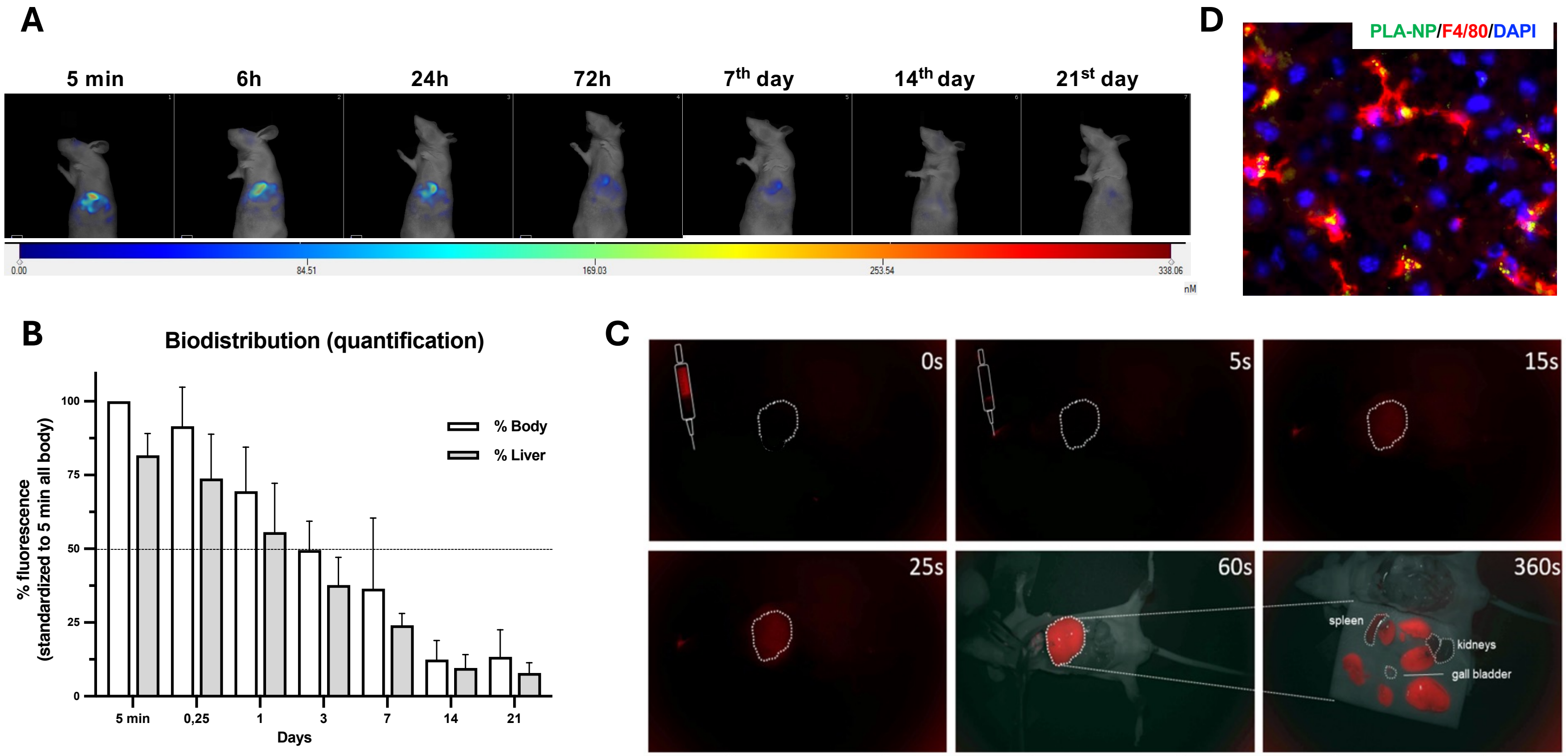

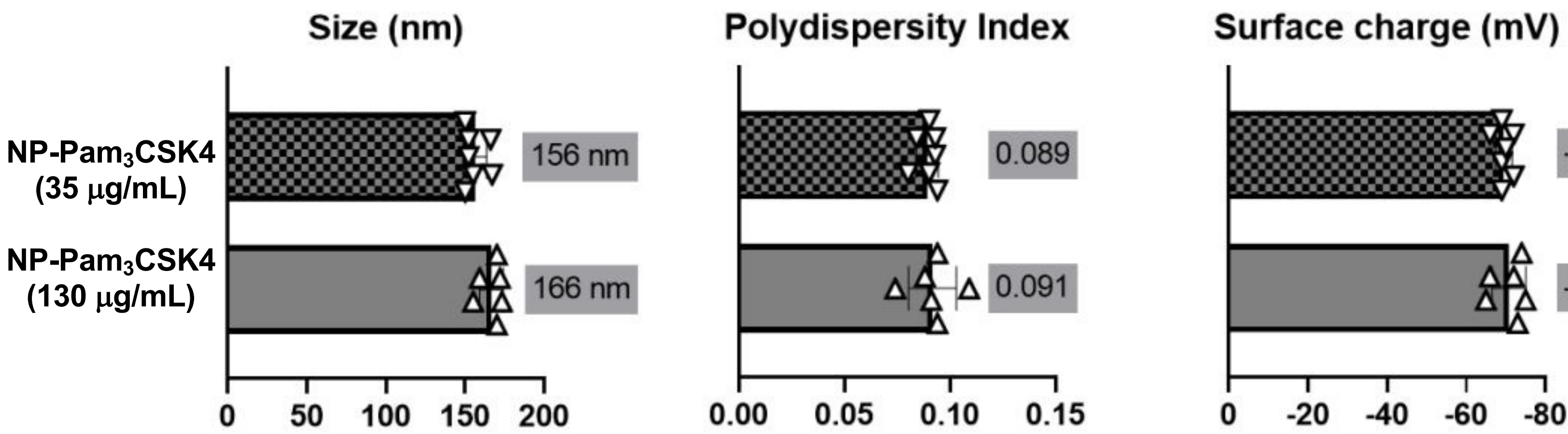

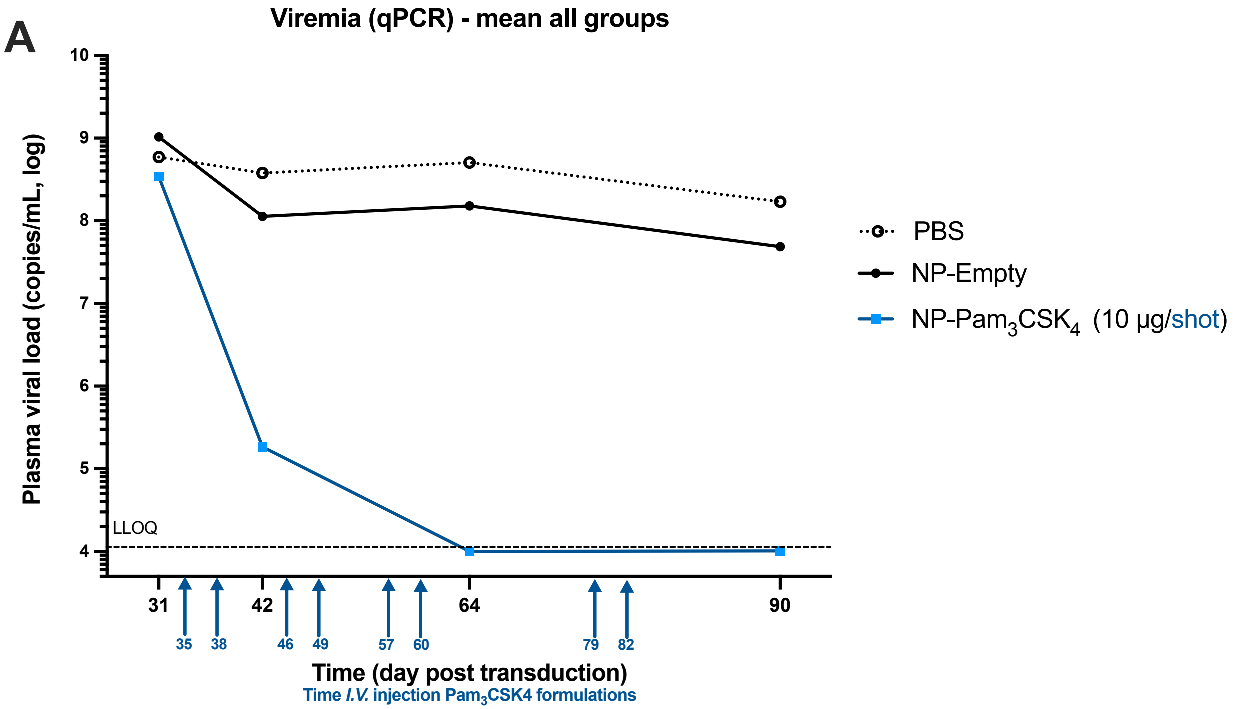

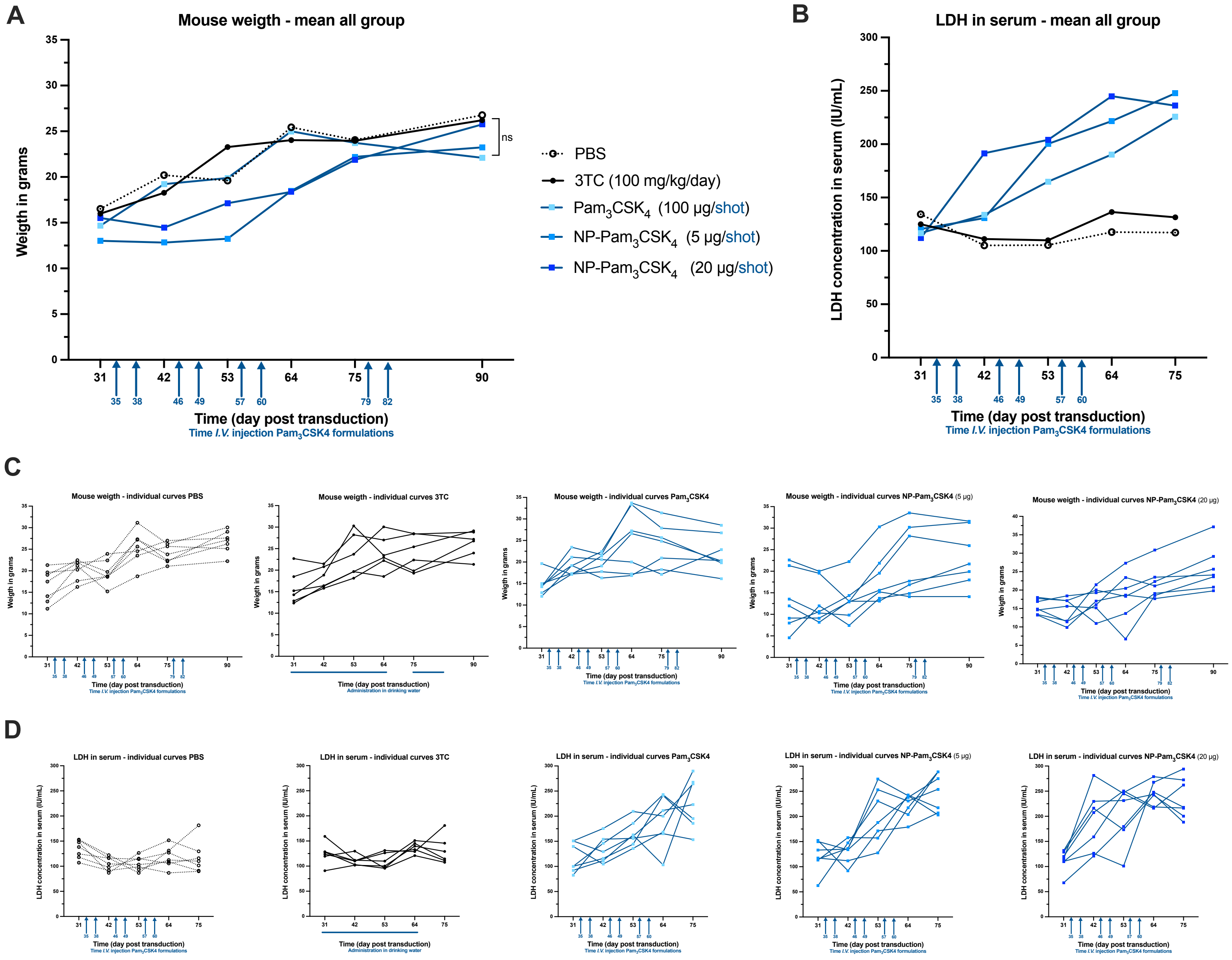

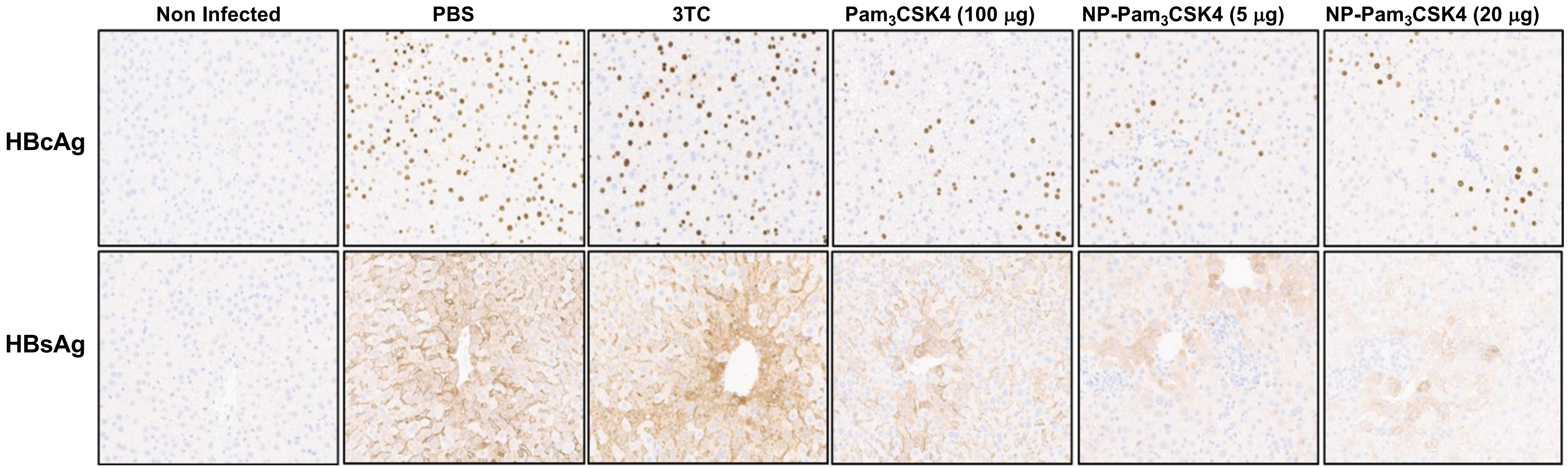

#### Figure S7

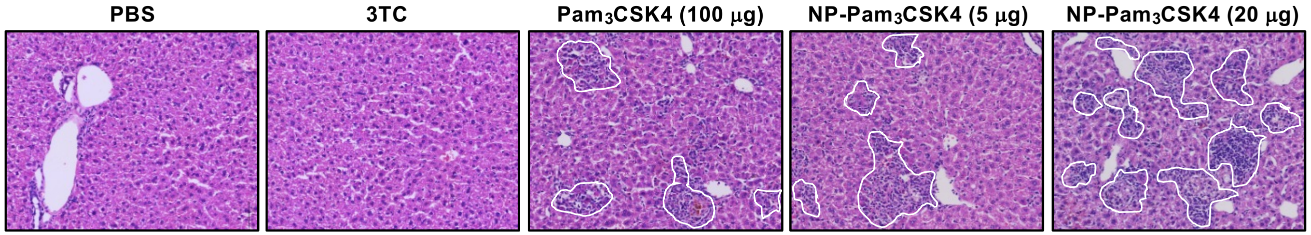

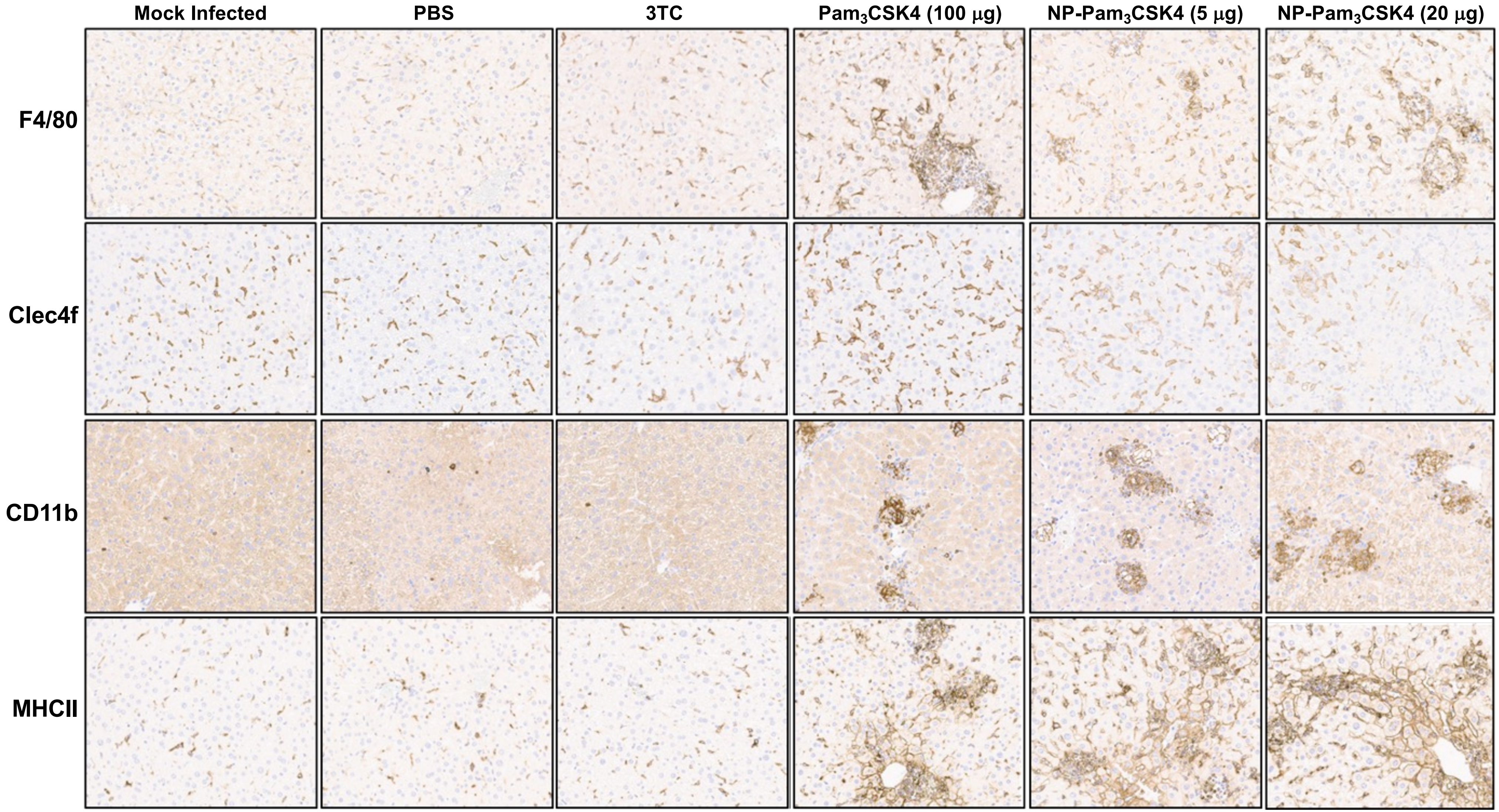

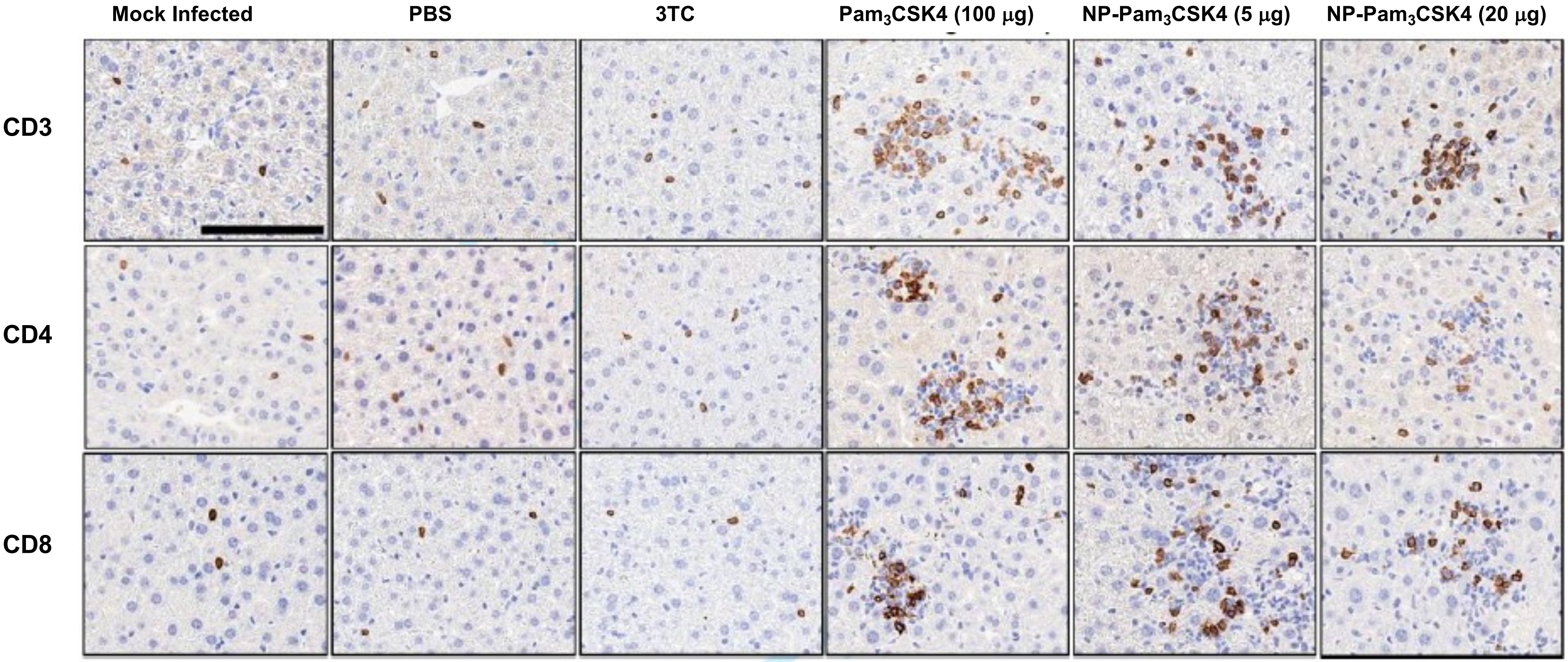
